# Rapid propagation of membrane tension at a presynaptic terminal

**DOI:** 10.1101/2021.05.26.445801

**Authors:** Carolina Gomis Perez, Natasha R. Dudzinski, Mason Rouches, Benjamin Machta, David Zenisek, Erdem Karatekin

**Author notes:** These authors contributed equally (Carolina Gomis Perez, Natasha R. Dudzinski).

## Abstract

Many cellular activities, such as cell migration^1^, cell division^2,3^, signaling^4,5^, infection^6^, phagocytosis^7^ and exo-endocytosis^8–11^, generate membrane tension gradients that in turn regulate them^12^. Moreover, membrane flows, which are driven by tension gradients, can limit exo-endocytosis coupling in space and time, as net membrane flow from exocytic to endocytic sites is required to maintain membrane homeostasis^13^. However, there is controversy over how rapidly plasma membrane flows can relax tension gradients; contrary to the common view^12,14,15^, recent work showed membrane tension does not equilibrate in several cell types^16^. Here we show membrane tension can propagate rapidly or slowly, spanning orders of magnitude in speed, depending on cell type. In a neuronal terminal specialized for rapid synaptic vesicle turnover and where exo-endocytosis events occur at distinct loci, membrane tension equilibrates within seconds. By contrast, membrane tension does not propagate in neuroendocrine adrenal chromaffin cells secreting catecholamines. Thus, slow membrane flow and tension equilibration may confine exo- and exocytosis to the same loci^17^. Stimulation of exocytosis causes a rapid, global decrease in the synaptic terminal membrane tension, which recovers slowly due to endocytosis. Our results demonstrate membrane tension propagates rapidly at neuronal terminals and varies during synaptic activity, likely contributing to exo-endocytosis coupling.

---

Neuronal presynaptic terminals are hubs of intense and rapid membrane turnover. Upon stimulation, the equivalent area of the synaptic vesicles (SVs) that fuse rapidly with the presynaptic plasma membrane is recovered through compensatory endocytosis to restore cell membrane area and to maintain a releasable pool of SVs (Figure 1a). How exocytosis triggers endocytosis is under debate^18,19^; a possible mechanism involves sensing at the endocytic site a drop in cell membrane tension upon exocytic membrane addition some distance away^20^, supported by the observation that even modest increases in membrane tension strongly inhibit endocytosis^21–23^. However, recent work indicated that membrane tension gradients in several cell types do not equilibrate over micron length scales for tens of minutes, implying extremely slow membrane flows^16^. Membrane tension propagating so slowly could not possibly act as a signal to trigger rapid compensatory endocytosis even at sites a few hundred nanometers away from typical release sites^20^. Even if a signal other than a decrease in membrane tension triggered endocytosis, membrane flow would be too slow to supply membrane to endocytic locations in the periphery of active zones, at least in terminals with rapid synaptic vesicle turnover^13^. These results suggest that either synaptic terminals must be specialized for facile membrane flow, or that sensing exocytosis and supplying membrane to endocytic sites must occur via other means.

**Figure 1.**
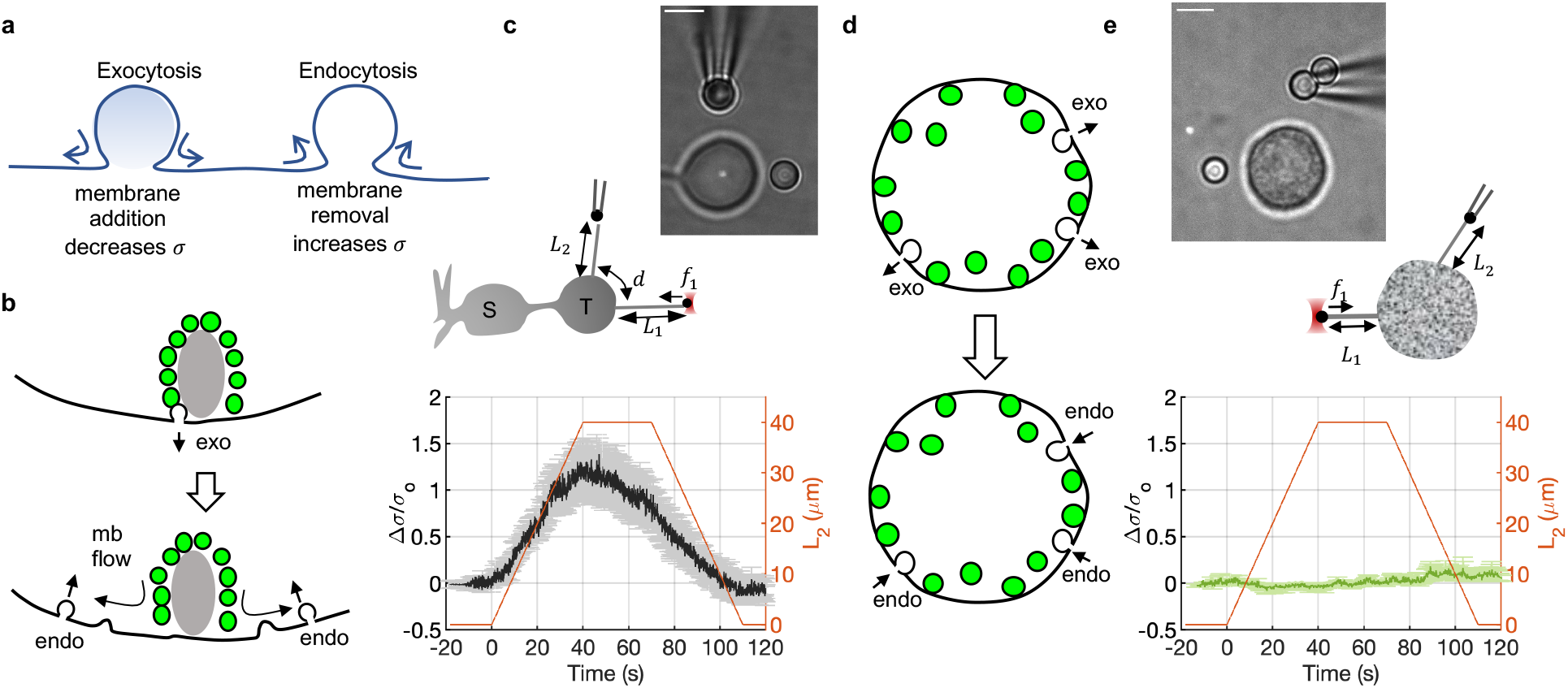
Membrane tension propagates rapidly in neuronal synaptic terminals, but not in endocrine chromaffin cells. **a**. Exocytosis adds membrane area to the plasma membrane, locally decreasing membrane tension, *σ*. Endocytosis locally increases *σ* through removal of plasma membrane area. If exo- and endocytosis occur at distinct loci, net membrane transfer from the exocytic to endocytic sites would be needed to sustain continuous vesicle recycling. **b**. In goldfish retinal bipolar terminals release predominantly occurs at active zones marked by ribbons where synaptic vesicles are tethered (depicted as a grey oval). Endocytosis sites are distributed throughout the terminal; thus most endocytosis occurs at loci distinct from active zones. **c**. Testing membrane flow at bipolar cell terminals. Middle. Schematic of the experiment. A “probe” tether is pulled from the terminal of a bipolar neuron using a 3 *μ*m diameter latex bead held in an optical trap (OT). A second, “pulling” tether is extended from the terminal using another bead held by a micropipette mounted on a 3-axis programmable piezoelectric stage. While the length *L_2_* of the pulling tether is extended at 1 *μm/s* by 40 *μm*, held for 30 *s*, then returned to its original position, the probe tether is held stationary and its tension is measured through the force acting on the bead held in the OT. S: soma, T: terminal, D: dendrites. Top: a snapshot from an experiment. Only the terminal and part of the axon of the cell are visible. Bottom: change in membrane tension Δ*σ* of the probe tether relative to its resting value, *σ*_0_, as a function of time. Average from 12 cells is shown. Error bars (gray) indicate standard error. Inter-tether distance *d* was 4 – 11 *μm*. The length *L_2_* of the pulling tether is shown on the right-axis (red). **d**. In adrenal chromaffin cells, most secretory granules undergoing exocytosis are retrieved at the same loci. **e**. Same as in **c**, but tethers were pulled from a neuroendocrine chromaffin cell. Average of 6 cells is shown. Inter-tether distance *d* was 6 – 12 *μm*. Scale bars: 5 μm.

Here we used goldfish retinal bipolar neurons as a model to test these possibilities, as they possess a single, giant (~10 μm) terminal, filled with up to 50 active zones marked by a ribbon, a dense structure to which approximately 100 SVs are attached and at the base of which most release takes place^24,25^. Each terminal contains a total pool of ~0.5 million SVs^26,27^. These cells release glutamate rapidly (time constants^28^ ~1-2 ms and 150 ms) to graded potentials: up to ~10-15% of the initial area can be added to the terminal within 250 ms upon strong stimulation^28–31^. Endocytosis restores terminal membrane area at longer time scales (~2 s and ~30 s for weak and strong stimuli^29^). Importantly, in these cells most endocytosis is known to occur away from ribbon sites where exocytosis occurs^32,33^, implying relatively fast membrane flow (Figure 1b). As a comparison, we tested membrane tension propagation and flows in neuroendocrine adrenal chromaffin cells. In contrast to neurons, these cells release their cargo in less spatially restricted regions into the blood stream, undergoing exocytosis and endocytosis in overlapping regions^17,34^ which does not necessitate long-range membrane flows. We find that membrane flow is remarkably rapid in bipolar cell terminals, while it is at least two orders of magnitude slower in adrenal chromaffin cells, suggesting that the dramatically different cell membrane hydrodynamics reflect unique functional needs of different cell types (Extended Data Figure 1).

## Membrane tension dynamics

To probe propagation of membrane tension in bipolar cell presynaptic terminals, we pulled a pair of thin membrane tethers from the cell surface using 3 μm latex beads as handles (Figure 1c). First, a short “probe tether” was pulled using an optical trap (OT) and maintained at constant length. Then a second, “pulling tether” was extruded a few μm using a bead manipulated by a micropipette mounted on a programmable piezo stage. After equilibration, we extended the pulling tether at 1 μm/s for 40 μm, held it for 30 s, then relaxed it to its initial position, while monitoring tension changes in the probe tether using OT. Extension of the pulling tether caused a local increase in cell membrane tension (Extended Data Figure 2). Propagation of this local change to the probe tether was monitored *via* the force acting on the probe tether using OT, 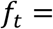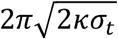, where *σ_t_* is the tether membrane tension and κ ≈ 0.27 pN·μm is the bending modulus^35^. The tether force is deduced from deviations of the bead’s position from the center of the OT and known calibration of the trap stiffness, allowing us to estimate *σ_t_* (Methods). To prevent spontaneous depolarizations which lead to exo-endocytosis^36^ that could perturb membrane tension measurements, we maintained cells in a low extracellular calcium solution.

Remarkably, membrane tension perturbations created by the pulling tether were transmitted rapidly, within at most a few seconds, to the probe tether, despite being separated by 4 – 11 μm (Figure 1c, Supplementary Movie 1). As the pulling tether was extended, increasing tension in the tether and its base (Extended Data Figure 2), tension in the probe tether increased. When extension of the pulling tether stopped, tension in the probe tether started relaxing. When the length of the pulling tether was decreased, tension in the probe tether decreased (Figure 1c).

Additional experiments confirmed these observations and ruled out potential artifacts. First, using similar double-tether experiments, we confirmed that in HeLa cells membrane tension does not propagate over micrometer length scales within several minutes, consistent with a recent report^16^ (data not shown). Second, we confirmed that tethers pulled from bipolar cells are cytoskeleton-free (Extended Data Figure 3), as the force-tension relationship above assumes. Third, we confirmed that membrane tension in the probe tether tracked tension changes in the pulling tether by monitoring fluorescence changes simultaneously in the two tethers (Extended Data Figure 4). We conclude that membrane tension equilibrates over several microns within a few seconds in presynaptic terminals, in stark contrast to other cell types^16^.

To probe propagation of membrane tension in the soma, we repeated the double tether experiments of Figure 1c but pulled both tethers from the soma, with inter-tether distances 4-12 μm (n=8). We found the membrane tension perturbation created by the pulling tether was again transmitted rapidly to the probe tether, but the change in membrane tension relative to its initial value was smaller at the probe tether (Extended Data Fig. 5). These results were consistent with fluorescence-based simultaneous tension estimates of the two tethers (Extended Data Fig. 4e). The resting tensions covered a broad range and were indistinguishable in the somata and terminals, while the tension perturbation created by extending a tether increased in a similar manner (Extended Data Figure 2c-e). Thus, membrane tension also propagates rapidly in the soma. Terminal-to-axon transmission of membrane tension changes were also rapid, with a slightly lower amplitude, but inter-tether distances were larger (14-17 μm, n=4, Extended Data Figure 5).

As a comparison, we tested how rapidly tension propagates in neuroendocrine chromaffin cells, where endo- and exocytosis occur in overlapping regions, using double-tether experiments as above. Extending the pulling tether strongly increased membrane tension locally (Extended Data Figure 2), yet, we could not detect membrane tension propagation over 6-12 μm in several minutes in any of the 6 cells tested (Figure 1e and Supplementary Movie 2).

We conclude that membrane tension propagates very rapidly in bipolar cells and unmeasurably slowly in adrenal chromaffin cells, two cell types specialized for secretion but with different spatiotemporal secretory vesicle dynamics, the latter not requiring long-range plasma membrane transport.

## Tracer diffusion and immobile obstacles

We tested whether a difference in the mobility of integral transmembrane domain (TMD) proteins could explain the large differences in membrane flows we observed among chromaffin cells, bipolar neuronal terminals, and somata. We labeled surface proteins with an organic dye and used fluorescence recovery after photobleaching (FRAP), fitting measured curves to find an immobile fraction and a diffusion constant of mobile labeled proteins (Extended Data Figure 6). Surprisingly, we found no significant difference among the tracer diffusivities for the three cases (*D_t_* = (10 ± 6.7) × 10^−3^, (19 ± 5.8) × 10^−3^, and (11 ± 4.8) × 10^−3^ μm^2^/s, for chromaffin cells, termini, and somata, respectively). The immobile fraction was similar for the terminal (0.39 ± 0.02) and soma (0.37 ± 0.04), which were ~30% lower than for chromaffin cells (0.60 ± 0.03).

We wanted to understand whether the differences in TMD protein immobile fractions could explain the differences in membrane tension propagation. The presence of immobile obstacles can dramatically hinder bulk flow even while tracer diffusion remains relatively unimpeded, especially in 2-dimensions (2D)^16,37^. We tested if a recent model^16^ in which TMD proteins that interact with the underlying cytoskeleton represent a random array of fixed obstacles to flow could explain our observations. The model predicts diffusive membrane tension propagation, with a tension diffusion coefficient *D_σ_* = *Ek/η*, where *E* is the membrane stretch modulus, *η* is the 2D membrane viscosity, and *k* is the Darcy permeability of the array of obstacles. In a simple model where obstacles are randomly arranged in space, the Darcy permeability is a function of the obstacle area fraction *ϕ* and the radius *a* of obstacles^16,37^, *k* = *a*^2^*f*(*ϕ*), where *f*(*ϕ*) is a rapidly decaying function with increasing *ϕ*. We estimated *ϕ* from the immobile membrane protein fraction we measured above. Assuming ~25% of membrane area is occupied by transmembrane proteins^38,39^, *ϕ* ≈ 0.093 ± 0.01, *ϕ* ≈ 0.096 ± 0.005 in the soma and terminal, respectively, and *ϕ* ≈ 0.149 ± 0.007 in chromaffin cells, yielding *k* ≈ 7.45 ± 3.7 nm^2^ and 6.96 ± 2.3 nm^2^ for the soma and terminal, and *k* ≈ 3.01 ± 1.06.7 nm^2^ for chromaffin cells. Assuming similar values for *a*, *η* and *E* for bipolar and chromaffin cells, only a 2.4-fold difference in the tension diffusion coefficient *D_σ_* would be expected, which is insufficient to explain the differences of membrane tension propagation between chromaffin cells and bipolar neurons. Using numerical calculations, we estimate 100-1000-fold larger *D_σ_* is needed to explain the rapid membrane flow we observe in bipolar cells compared to chromaffin or HeLa cells, if tension propagates diffusively (Figure 2a, see also Supplementary Information and Extended Data Figure 7). In addition, the speed and amplitude of tension propagation show little, if any, distance dependence (Figure 2b,c), suggesting the underlying propagation mechanism may not be diffusive in bipolar neurons. In summary, neither tracer diffusion nor the fraction of immobile obstacles can explain the large differences in membrane tension propagation we observe in bipolar and chromaffin cells. Thus, while our results cannot rule out a model in which the cell membrane flows are heavily disturbed by immobile TMD proteins^16^, other factors likely contribute to the propagation of membrane tension in bipolar cells.

**Figure 2.**
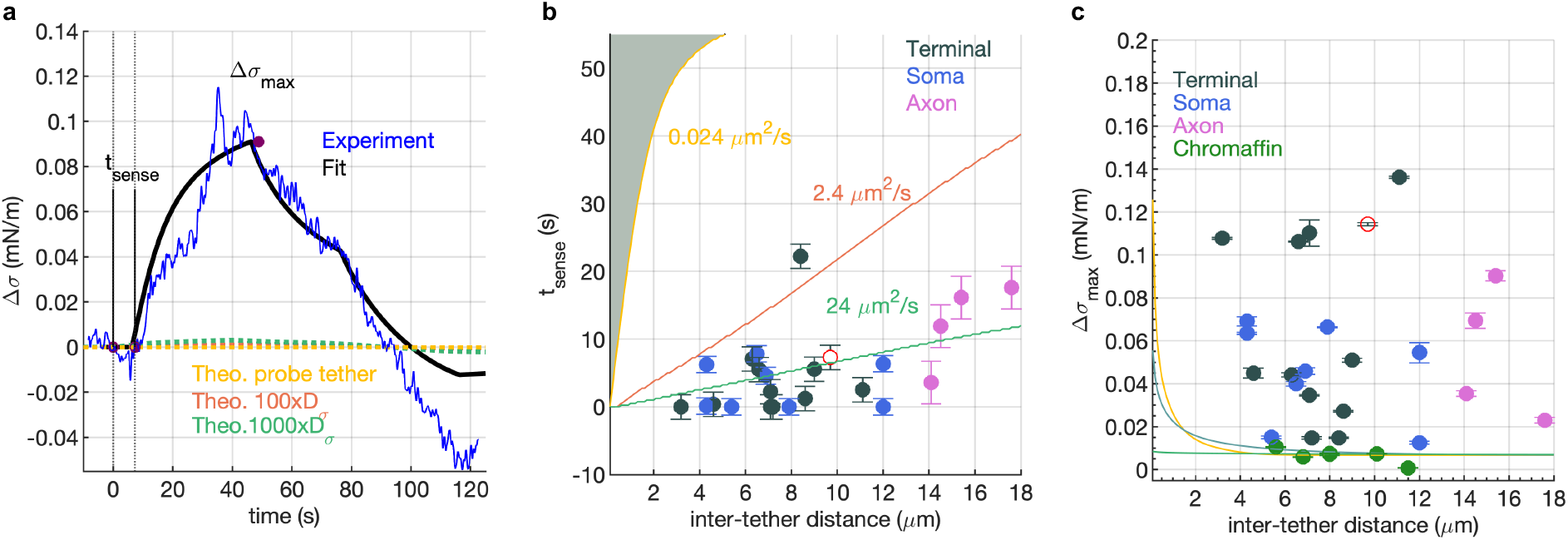
Rapid membrane tension propagation in bipolar cell terminals is unlikely to be diffusive. **a.** Comparison of measured and predicted membrane tension changes at the probe tether as the pulling tether creates a controlled tension perturbation a distance *d* away (see Figure 1). A model of membrane flow through a random array of immobile obstacles predicting diffusive propagation of membrane tension^*16*^ was fitted to experimentally measured membrane tension changes at the probe tether. In this example, the inter-tether distance was *d* = 9.7 μm. Membrane tension diffusion coefficients varying from the value estimated^*16*^ for Hela cells (*D_σ_* = 0.024 μm^2^/s) to a value 10^3^-fold larger did not produce a good fit. Good agreement was obtained only when *d* was assumed to be 0.1 μm (black) suggesting very rapid propagation of tension with weak distance dependence. **b**. The time to sense at the probe tether a tension change at the pulling tether, *t_sense_*, as a function of inter-tether distance *d*. To estimate *t_sense_*, we used the difference between the actual time the pulling tether was set in motion (*t* = 0) and the onset of the predicted best fit tension change at the pulling tether (black curve in **a**). Filled circles are experimental measurements. The red empty circle corresponds to the trace in **a.** The values of *t_sense_* predicted by diffusive propagation of membrane tension are indicated as solid curves, with the corresponding values of *D_σ_*. The hatched region corresponds to values of *t_sense_* allowed by the model and measurements of Shi et al.^*16*^. **c.** Maximum change in Δ*σ* at the probe tether as a function of distance from the pulling tether. The amplitude-distance relationship predicted by the diffusive propagation model is indicated by the solid curves (same color scheme as in a,b). Filled circles are the experimental values and the empty red circle corresponds to the trace in **a**. (see Supporting Information and Extended Data Figure 7 for details, parameter values and all fits).

## Plasma membrane-cytoskeleton drag

The dominant resistance to membrane flow in cell membranes arises from interactions of the plasma membrane with the adjacent layer of the underlying cytoskeleton^16,40^. Forcing membrane tethers to slide on the cell surface and monitoring the force-velocity relationship provides a measure of this resistance^41^.

We found most tethers (12/15) could be dragged with ease around the bipolar cell terminal as soon as a finite tangential force was generated by moving the cell with respect to the bead holding the end of the tether (Figure 3a,b, and Supplementary Movie 3). Occasionally, tethers were transiently immobilized for 1-3 s. Combining image analysis with force measurements, we quantified, *f*_∥_, the force tangent to the terminal membrane and the distance traveled by the tether base for every frame (Figure 3c). From the distance profile, we computed the frame-to-frame velocities. Tether base velocity as a function of tangential force is shown in Figure 3d (black circles). Low forces (~6 pN) were sufficient to produce large velocities. Tethers moved mostly unimpeded, interspersed with transient pauses, likely due to the heterogeneous nature of the underlying cytoskeleton. By contrast, most tethers extruded from chromaffin cells (8/10) were immobile. Tethers that did slide here required much larger forces (~24 pN), and paused more frequently after initial sliding for ~0.25-1 μm (Figure 3h-j and Supplementary Movie 4). Tethers dragged from bipolar cell somata displayed intermediate behaviour (Figure 3e-g, Extended Data Figure 8 and Supplementary Movie 5).

**Figure 3.**
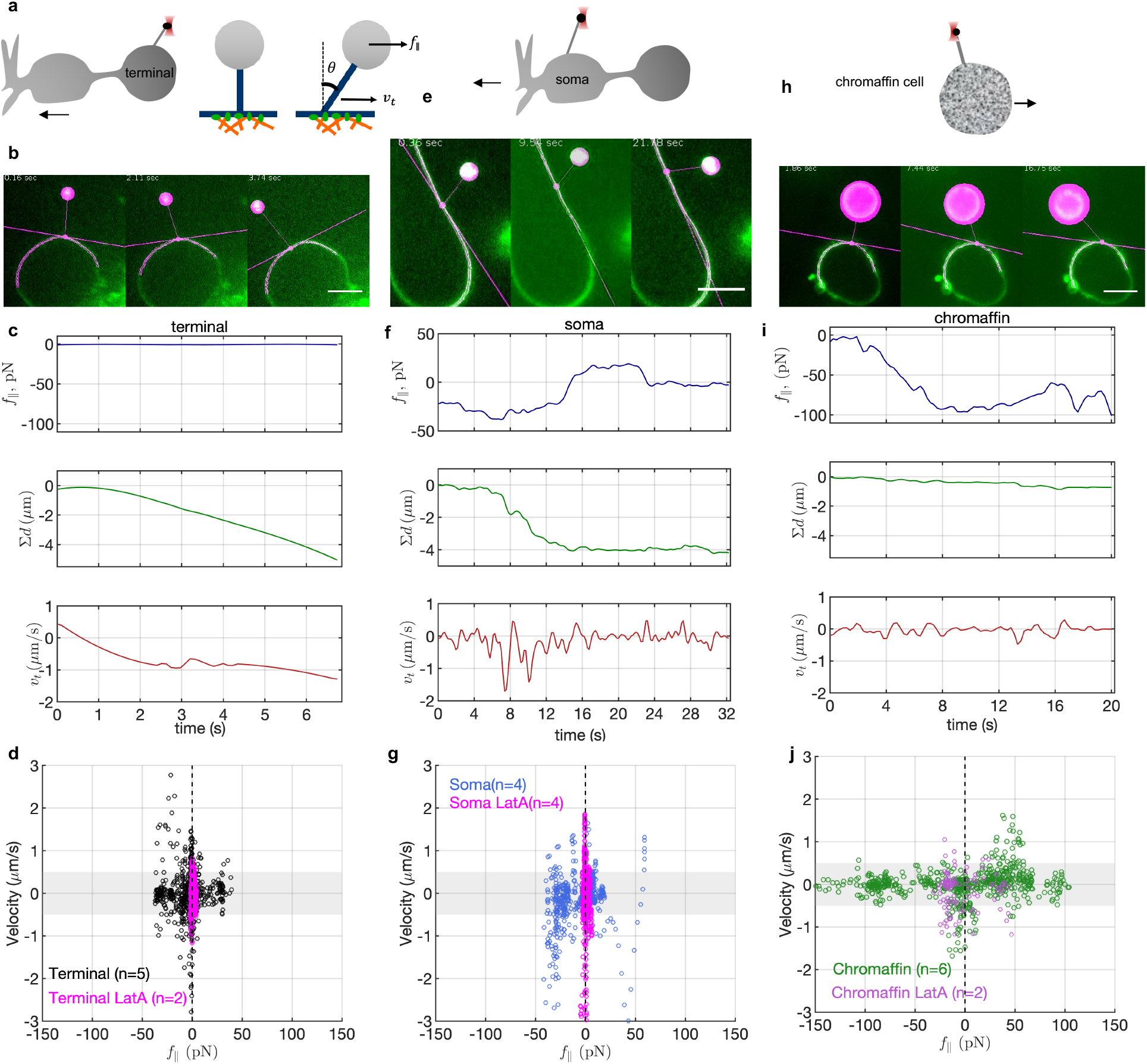
Membrane tethers extruded from bipolar neurons, but not from chromaffin cells, can be dragged with ease. **a**. Schematic of the experiment. After a membrane tether is extruded from a terminal, the cell is moved to generate a tangential component *f*_∥_ of the tether force, given by the sine of the tether-membrane angle *θ*. The velocity *v_t_* with which the tether base moves is recorded. **b**. Snapshots from a tether-dragging experiment at a bipolar cell terminal. The detected contour of the cell membrane is shown in magenta. The intersection of the tether with the cell surface is indicated by a magenta point, and the tangent at that point by a line. The bead is automatically detected and overlaid by a disc. Cell membranes were visualized with CellMask Deep Red or FM4-64. **c**. Time profiles of the tangential force, *f*_∥_, cumulative distance traveled from origin, ∑ *d*, and frame-to-frame tether velocity, *v_t_*. Scale bar = 5 μm. **d**. Tether velocity-tangential force pairs for untreated (black) or LatA treated (20 μM, 20 min, magenta) terminals. The gray area, |*v_t_*| < 0.5 μm/s, corresponds to the level of noise in *v_t_* and may be considered stationary. **e-g**. Similar measurements for tethers drawn from the somata of bipolar neurons. **h-j**. Similar measurements for tethers drawn from adrenal chromaffin cells. For bipolar cell terminals, and to a lesser extent for somata, very small forces were sufficient to set a tether in motion, whereas tethers drawn from chromaffin cells required much larger forces to slide. LatA treatment reduced forces in all cases, even though there was no visible blebbing under these conditions.

Disruption of the actin cortex using latrunculin A (LatA, prevents actin polymerization^42^) facilitated tether sliding in terminals and somata, and to a lesser extent in chromaffin cells (Figure 3g,j, Extended Data Figure 8). After treatment with 20 μM LatA for 20 min, which does not lead to visible bleb formation, most tethers drawn from somata (5/7, 71%) could be dragged with ease and supported forces (〈*f*_∥_〉 ≈ 1.6 pN) similar to those in treated terminals (〈*f*_∥_〉 ≈ 0.9 pN). Tethers from treated chromaffin cells slid readily (2/2), albeit at high angles and with higher forces (〈*f*_∥_〉 ≈ 18 pN) than untreated termini or somata.

Together, these results suggest that the cytoskeleton impedes tether sliding to a much higher degree in chromaffin cells than in bipolar cell somata. Tethers drawn from terminals display minimal resistance to tether dragging, despite possessing a readily detectable F-actin cortex (refs. 43,44 and Extended Data Figure 3).

## Activity-dependent changes in membrane tension

Membrane area changes of the terminal have been monitored directly using time-resolved capacitance^29,30,45^: the excess area added to the terminal upon mild stimulation (up to 0.2 s depolarization to 0 mV, leading to fusion of ~2000 SVs, or ~8% of the terminal area) is recovered through rapid endocytosis (time constant ~1 s), whereas membrane area increases elicited by stronger stimulation are restored over ~20-30 s. If synaptic vesicles fuse completely with the active zone membrane^32,33^ and tension perturbations propagate rapidly in the terminal, we expect to observe a rapid, terminal-wide decrease in membrane tension upon strong stimulation of exocytosis, followed by slower recovery due to endocytosis.

To test this hypothesis, we used two approaches. First, we took advantage of the fact that some bipolar cells spontaneously develop calcium action potentials that drive exocytosis^36^. We loaded cells with a calcium indicator, Fluo-4-AM, to monitor calcium fluctuations^46^ while simultaneously measuring membrane tension at the terminal using OT. Occasionally, cells displayed robust calcium activity. Calcium spikes anti-correleated with membrane tension (Extended Data Figure 9), consistent with calcium-stimulated exocytosis reducing membrane tension and endocytosis driving its recovery^36^. Second, to exert better control over the release process, we used a photochemical approach to stimulate exo- and endocytosis (Figure 4). Neurotransmitter release in type Mb bipolar cells is driven by calcium entry via L-type calcium channels^47,48^. To stimulate release, we bathed the cells in nifedipine (20 μM), a blocker of L-type DHP-sensitive calcium channels^49^. Photolysis of nifedipine at 405 nm removes the block, allowing calcium influx and exocytosis^50^. We loaded cells with a calcium indicator, Fluo-4-AM, to monitor calcium entry upon stimulation^46^. We pulled a membrane tether and monitored its tension while imaging calcium signals using time-lapse spinning-disc confocal microscopy. Stimulation with 405 nm illumination for 1 s caused a rapid increase in the calcium signal (Figure 4b,c). There was a concomitant decrease in membrane tension with a characteristic time of ~1-2 s. Even though the cytosolic calcium increase and the accompanying exocytosis occur instantly at the time resolution of these recordings (1 frame/s), the tension decrease is slower, limited by how rapidly the cell membrane flows over the cell’s surface and into the tether (Extended Data Fig 2f). Membrane tension recovered more slowly, over a 10-20 s time scale (Figure 4c). The initial rapid decrease in membrane tension was likely due to exocytosis, as no change in cytosolic calcium or membrane tension occurred without nifedipine photolysis or when photolyis was carried out in the absence of extracellular calcium (Extended Data Figure 9e). The slow recovery of membrane tension was likely due to endocytosis, because treatment with myristyl trimethyl ammonium bromide (MiTMAB), an agent that blocks endocytosis by inhibiting dynamins I and II^51,52^, inhibited recovery. The recovered fraction of membrane tension within 50 s of stimulation was ~1.0, 0.49, and 0.29 with 0, 10, or 30 μM MiTMAB, respectively (Figure 4 and Extended Data Figure 9). We used pro-myristic acid, rapidly broken into myristic acid in the cytosol, as a negative control, which did not affect membrane tension dynamics (Extended Data Figure 9). No tension changes could be detected in the soma during neuronal activity (Figure 4d). Thus, synaptic vesicle recycling generates large changes in membrane tension which are mostly confined to the terminal.

**Figure 4.**
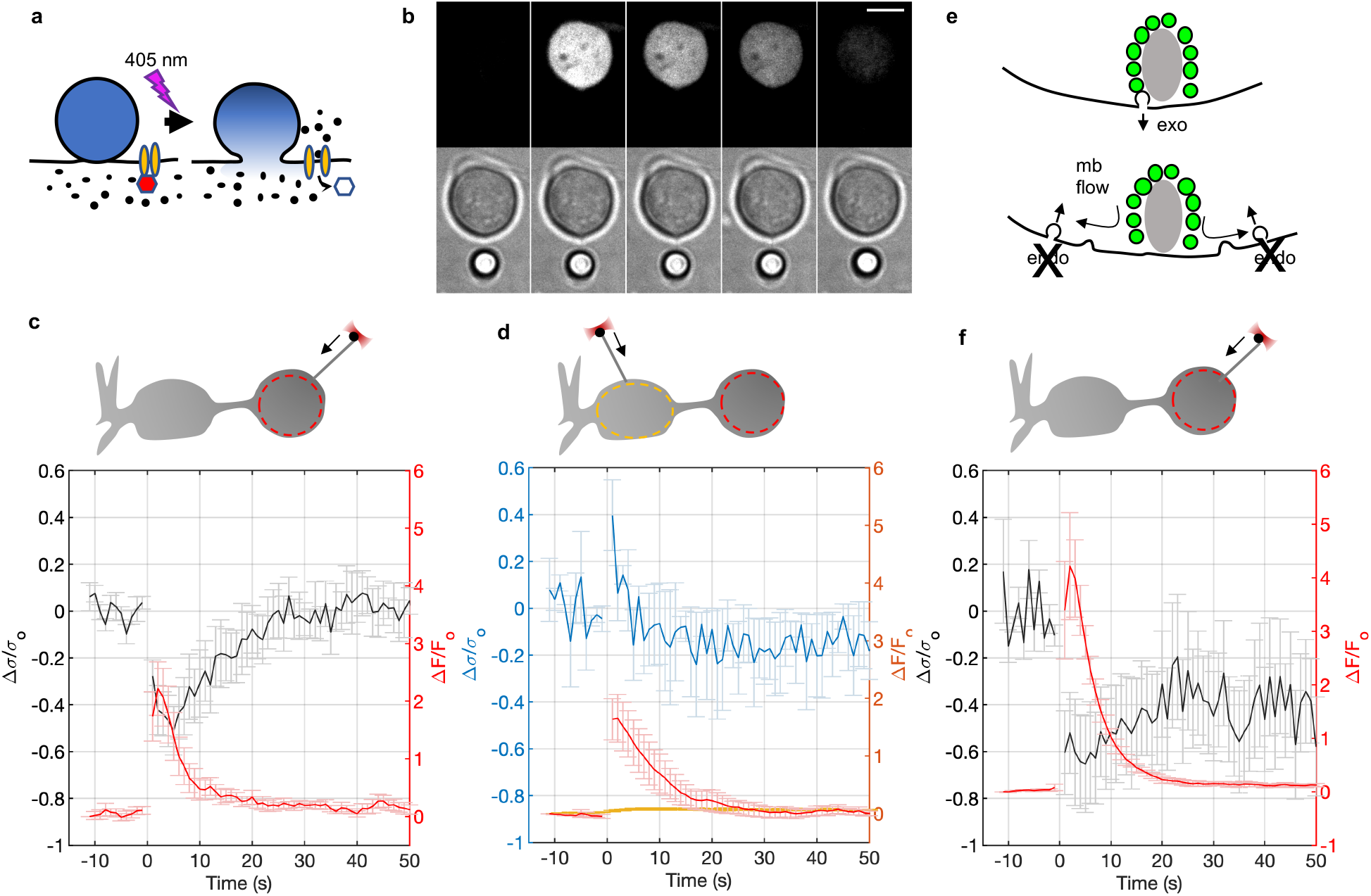
Activity-dependent changes in membrane tension in bipolar neurons. **a**. Principle of photstimulation employed. Nifedipine (depicted as a red hexagon) blocks L-type calcium channels. Photolysis at 405 nm removes the block, causing calcium entry and exocytosis^50^. **b**. Example of a simultaneous calcium-imaging and tether-force measurement during photostimulation of a bipolar cell terminal. A membrane tether was pulled from a bipolar cell terminal and membrane tension monitored while nifedipine in the bath (20 μM) was photolysed by a 1 s 405 nm pulse. The cell was pre-loaded with the calcium indicator Fluo-4. Top row: Fluo-4 fluorescence. Bottom row: brightfield imaging. The two channels were alternated during acquisition. The tether becomes visible upon stimulation and calcium increase (2nd column), indicating a sharp decrease in tether tension. Scale bar = 5 μm. **c**. Fractional changes in membrane tension (black, left axis) and calcium signals (red, right axis) averaged over 6 experiments. Photolysis was applied for 1 s starting at *t* = 0 s. Membrane tension recovered completely within 50 s after stimulation. The region of interest used for fluorescence quantification is indicated in the schematic as a dashed circle. **d.** Similar to **c**, but the tether-forces were measured in the soma (n=4). Most L-type calcium channels are in the terminal where calcium dynamics are largely confined^48^. **e**. Inhibition of endocytosis by MiTMAB, a dynamin inhibitor^51,52^, is expected to reduce recovery of membrane tension following exocytosis. **f.** Similar to **c**, but measurements were made in the presence of 30 μM MiTMAB to block endocytosis (n=4). Only ~30% of the initial drop in membrane tension upon stimulation recovered. See Extended Data Figure 9 for additional experiments.

## DISCUSSION

We have shown that cell membranes can flow at vastly different speeds, likely reflecting physiological requirements. In neuronal presynaptic terminals, rapid membrane flow is likely an adaptation required for maintaining SV exo-endocytosis at distinct loci, as recognized nearly 50 years ago^13^. Because endocytosis is inhibited strongly with increased membrane tension^21–23^, fast terminal-wide equilibration of membrane tension gradients likely contributes to the signaling that couples exo- and endocytosis^20,29^. Conversely, slow membrane flow likely limits the spatial and temporal extent of exo-endocytic coupling in neuroendocrine adrenal cells. Upon fusion, secretory granules rarely collapse into the plasma membrane in such cells^17,34^, and those that do take seconds to do so^17,53^, consistent with the extremely slow membrane flow we observed in chromaffin cells.

Resistance to membrane flow primarily arises from interactions of the cell membrane with the underlying cytoskeleton^12,16,40^. Consistent with hydrodynamic models of 2D flow around fixed obstacles^16^, we found tracer mobility is a poor predictor of membrane flow. The fraction of immobile membrane proteins is a better indicator, but cannot quantitatively explain the large differences between membrane flows we observed. Tether drag and its sensitivity to agents disrupting the cortical F-actin network correlates much better with the observed membrane flows. Thus, it is likely that membrane tension dynamics are governed not only by the density of the immobile obstacles, but also by their spatial arrangements^54,55^, hydrodynamic drag between the membrane and the cytoskeleton, and how easily the immobile obstacles can be detached from the cytoskeleton under flow^40,56^. Like chromaffin cells^57^, bipolar cell terminals possess a dynamic F-actin cortex^43,44^, but the arrangement of the actin cytoskeleton, how it is linked to the PM, and how these interactions affect membrane flows at presynaptic terminals will need to be addressed in the future.

## Supporting information

Supplementary Discussion

SI Guide, captions of SI Movies

## METHODS

### Cell and tissue preparation

All procedures for animal care were carried out according to Yale Animal Care and Use Committee (YACUC). Goldfish retinal bipolar neuron dissection and dissociation was carried out following ref. 58. Goldfish were first dark adapted for at least 20 min., then decapitated using a scalpel and promptly pithed. Eyes were removed, hemisected, and placed in oxygenated dissociation buffer containing (in mM) 120 NaCl, 2.5 KCl, 0.5 CaCl_2_, 1 MgCl_2_, 10 HEPES, 0.75 EGTA, 10 Glucose, pH 7.4, 260 mOsm (osmolarity was checked with a Precision Systems Micro-Osmette Osmometer). Retina was removed from eyecups and placed in hyaluronidase solution 1100 units/ml (Sigma, H3884), dissolved in dissociation buffer to remove vitreous for a minimum of 12 minutes. Retinas were removed from hyaluronidase solution and cut into four to six pieces (each about 3 mm x 3 mm). Retina pieces were digested with papain solution (12.5 mg/ml, Sigma 76220) for 30 minutes containing 0.5 mg/ml L-cysteine (Sigma C7352) added to dissociation buffer. Pieces of tissues were removed from papain, rinsed in dissociation buffer, and kept at 12-14°C for 4-6 hours. To obtain dissociated retinal bipolar neurons, retina pieces were mechanically triturated using a fire polished Pasteur pipette and plated onto glass-bottom MatTek dishes coated with Poly-d-lysine (MatTek, P35GC-1.5-14-C). Retinal bipolar neurons were visually identified by morphology.

Mouse chromaffin cells from 1-month-old mice (C57Bl6) were cultured following ref. 59. Briefly, animals were euthanized using isoflurane overdose followed by cervical dislocation, the abdomen was open and both adrenal glands were removed. The medullas were dissected using a stereo microscope and transferred to a sterile Petri dish containing 1 mL of sterile ice‐cold Locke’s solution containing (in mM) 154 NaCl, 5 KCl, 3.6 NaHCO_3_, HEPES, and 11glucose. Tissues were disaggregated using papain (60–90 UI/mL) for 15–20 min at 37°C without shaking. Subsequently, tissues were washed with 800 μL of fresh Locke’s solution and passed through 1 mL and 100 μL pipette tips, until the suspension became turbid. Cells were plated onto glass-bottom MatTek dishes coated with Poly-d-lysine.

### Optical tweezers

The setup consists of a Perkin-Elmer Ultraview spinning-disc confocal system with a NikonTE2000 inverted microscope, Yokogawa CSU-X1 scanhead, a Hamamatsu C9100-50 EMCDD camera, and laser lines for 405, 488, 532, and 640 nm, controlled by Volocity or μ-manager software^60^. An optical trap is generated by focussing a 1064 nm laser beam (Coherent Matrix, 10 W) through a 100x/1.45 oil objective lens (Nikon Plan Apo λ) ~10 μm above the coverslip surface. Samples are moved by a piezoelectric stage (300 μm x 300 μm range, Piezoconcept, France), which is controllable via a joystick, Labview Virtual Instruments (VIs), or analog signals from a waveform generator (Agilent 33522A). A micropipette holder is attached to another, programmable 3-axis piezo unit (100 μm range per axis, P-611.3 NanoCube, with controller E-727 and Mikromove software, Physik Instrumente, Germany) that is used for programmable motion of the “pulling tether”. The 3-axis piezo stage is mounted onto a Newport manual M-462 Series xyz stage for coarse movement.

Trap stiffness was calibrated using a hydrodynamic flow method, following ref. ^61^. In a closed sample cell, the stage is oscillated with a sine wave, with peak-to-peak amplitude *A_pp_* and drive frequency *f_d_*, while a bead is trapped with OT. The bead positions are recorded using the EMCCD camera, at rates up to 85 Hz from small region of interest around the bead. The stage velocity *v_d_* is assumed to be equal to the liquid velocity, *v_l_*. Inertial forces are negligible because the Reynolds number is typically small: *R_e_* = (*v_l_* × *D_bead_* × ρ)/η~10^−6^ with *D_bead_*~1 μm, ρ~10^−9^ *pN* · *s*^2^/μ*m*^4^ (1 *kg/m*^3^), η~10^−3^ *pN* · *s*/μ*m*^2^, *v_l_*~1 − 10 μ*m*/*s*. The bead experiences a hydrodynamic force *f_hydro_* = γ(*v_b_* − *v_l_*), where *v_b_* is the bead velocity and γ = 6*πηR_bead_*. In the regime *w_d_* · τ ≪ 1, where *w_d_* is the angular frequency of stage oscillation (*w_d_* = 2π*f_d_*) and τ = γ/*k_Trap_*, the amplitude of the bead’s motion is *A_bead_* = *τw_d_A_pp_*/2. Thus, *f_bead_* = *k_Trap_ A_bead_* = *γw_d_A_pp_*/2 = *f_hydro_*. Trap stiffness *k_Trap_* is found from the slope of a linear fit to *A_bead_* vs *f_hydro_*. Trap stiffness was found to be linear with laser power. Trap stiffnesses used were in the range 73-271 pN/μm. We used carboxylated latex beads of 1.9 ± 0.16, or 3.2 ± 0.19 μm (mean±SD) diameter (Invitrogen, referred to in text as 2 or 3 μm beads).

For tether force measurements, a bead was trapped in the OT and its zero-force position, (*x_o_*, *y_o_*), was recorded for at least 10 s and averaged at rate ~14 or 30 frames/s. The bead was then brought into contact with the cell surface for ~1 s, and then pulled away from the membrane by joystick control of the piezo stage to form a membrane tether. Typically more than half of attempts were successful with bipolar cells but with chromaffin cells the success rate was somewhat lower. Bead positions (*x*, *y*) were recorded using digital image stacks. The presence of a tether was confirmed by visual assessment when possible (e.g. when the cell membrane was labeled with a dye or when the tether was visible in contrast-enhanced brightfield images) or by releasing the bead at the end of the experiment and observing it being pulled back to the cell surface. Bead positions were tracked offline using a custom-made MATLAB (Mathworks, MA) program that uses the Image Processing Toolbox function “imfindcircles” to detect the bead using a circular Hough transform algorithm, then calculates the weighted centroid of the bead. The deviation of the bead’s position 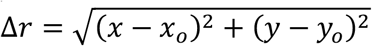 from its zero-force position was calculated for each frame and used to calculate the force acting on the bead, *f_b_* = *k_Trap_*Δ*r*, using the trap stiffness calculated as above. Imaging beads immobilized on a coverslip indicated the centroid of a bead can be detected with a root-mean-squared error of ~17 nm at 33 Hz sampling in brightfield images. With a typical *k_Trap_* = 75 pN/μm, this translates to an error in force estimation of *δf_b_* = 1.3 pN.

Membrane tension was estimated from the tether force acting on the bead^40,62^, 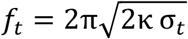, where *σ* is the membrane tension, and *κ* = 0.27 pN · μm is the membrane bending modulus for neuronal growth cones^62^, a value within the range reported for other cells^63–66^ (0.18 − 0.32 pN · μm).

### Double tether pulling

Goldfish retinal bipolar neurons were dissociated and plated into Ringers solution without Ca^2+^ (130 mM NaCl, 4 mM KCl, 1 mM MgCl_2_, 10 mM HEPES, 10 mM Glucose, 4 mM EGTA, pH 7.3, 260 mOsm). Chromaffin cells were plated into Locke’s solution lacking Ca^2+^. Calcium was omitted to prevent secretion that could affect membrane tension measurements. A micropipette with a ~1 μm diameter tip was attached to a pipette holder and lowered into the solution while applying positive pressure. Beads (3 μm diameter) were added to the dish, and a bead was caught at the tip of the pipette by applying negative pressure. Another bead was captured with the OT, and a tether (the “probe tether”) was pulled to a few μm length and held stationary. The bead held by the micropipette was then lowered further and used to pull a second tether (the “pulling tether”) from the cell surface to a few μm length using a joystick. After equilibration, the pulling bead was extended by 40 μm at 1 μm/s, held for 30 s, then returned to the initial position at 1 μm/s, using an automated protocol. Images were acquired at a rate of ~13-14 or 30 frames/s, except for experiments shown in Extended Data Figure 4 that used CellMask Deep Red fluorescence to track membrane tension changes, where the frame rate was limited to ~ 4.2 frame/s due to photobleaching.

### Photostimulation

For photostimulation, goldfish retinal bipolar neurons were dissociated and plated into a glass-bottom MatTek dish using 2 mM Ca^2+^ Ringers solution (125 mM NaCl, 4 mM KCl, 1 mM MgCl_2_, 2 mM CaCl_2_ 10 mM HEPES, 10 mM Glucose, pH 7.3, 260 mOsm) supplemented by 1μM Fluo-4-AM, a fluorescent calcium indicator dye, and 20 μM Nifedipine, a photolyzable L-type calcium-channel blocker. While allowing 30 minutes for attachment and dye uptake, a bipolar cell was visually identified. Immediately after washing with 2 mM Ca^2+^ Ringers solution without nifedipine or Fluo-4-AM, a membrane tether was pulled from the presynaptic terminal using a 3 μm diameter carboxylated latex bead. Images were recorded alternating between brightfield (to record the bead position) and fluorescence (488 nm excitation, 0.5% laser power, Chroma 527/55 nm band-pass emission filter, for calcium imaging using Fluo-4) at a rate 1 frame/s/channel. For stimulating calcium entry and secretion, nifedipine was photolysed using the 405 nm laser at full power for 1 s. Bead displacements were analyzed to determine membrane tension as above, and calcium dynamics were analyzed by plotting the fluorescence (background-corrected pixel values in a region of intetest) relative to its initial value, *ΔF/F_o_*.

### Tether Sliding

Goldfish retinal bipolar neurons were dissociated and plated into Ringers solution without Ca^2+^ (120 mM NaCl, 4 mM KCl, 1 mM MgCl_2_, 10 mM HEPES, 10 mM Glucose, 4 mM EGTA, pH 7.3, 260 mOsm) and with 5 μM CellMask Deep Red (Thermo Fisher) to visualize tethers. A few initial experiments used 14 μM FM4-64 (Invitrogen) instead of CellMask Deep Red, with no obvious difference in results. Tethers were extruded as above using a 2 or 3 μm diameter bead, either from the terminal or soma of a retinal bipolar neuron, or from a chromaffin cell. In some cases, 20 μM Latrunculin-A (LatA, Cayman Chemicals) was added to the bath during cell plating to disrupt the F-actin cytoskeleton.

#### Image Analysis

Tether sliding was quantified using a pipeline incorporating semi-automated image processing in ImageJ and analyses in MatLab. Bead force measurements were performed as described above, using a custom MatLab script that tracked the bead center through every frame utilizing the Computer Vision toolbox^67^. Videos were registered in the frame of the cell using ImageJ’s Linear Stack Alignment with SIFT plugin^68^. Significant registration drift was corrected with a manually tracked stationary landmark. The membrane contour was initiated manually for an initial frame, which was then tracked for that and subsequent frames using the Jfilament plugin^69^. The bead center was additionally tracked in the new reference frame. The point at which the bead contacts the membrane (the tether base) was manually tracked through each frame in ImageJ.

#### Data Analysis

Points from the tracked cell outline were fit with a spline, and the tangent line calculated at the nearest point to the tether base. The angle between the line perpendicular to the tangent and a straight line connecting the tether base to the bead defines the tether-membrane angle *θ*. The tangential force on the membrane *f*_∥_ is the sine of the tether-membrane angle *θ* multiplied by the total tether force at the given frame. The cumulative distance traveled by the tether base was extracted from each video. Cumulative distance and force traces were smoothed with a Gaussian with width 1 s. The tether velocity *v_t_* is then calculated per-frame as the derivative of the cumulative distance trace.

Forces and velocities were collated across individual experiments within each cell type for further analyses. The maximum supported force, *f_∥,max_* is the peak of the force trace when the tether is stalled, i.e. when velocity is below 500 nm/s, estimated to be the level of noise for a stationary tether. Obstacle density is the number of individual obstacles encountered divided by the total distance traveled by the tether base. Obstacles result in tether base stalling (*v_t_* < 500 nm/s) despite sustaining a substantial tangential force (*f*_∥_ > 5 pN). Obstacles were counted as distinct if separated by velocities *v_t_* > 500 nm/s.

### Tracer diffusion and immobile fraction of membrane proteins

Retinal bipolar neurons and chromaffin cells were labeled with 250 μg/ml Alexa-488-NHS dye (Thermo Fisher) for 30 min at room temperature (RT) or 37 °C respectively. Photobleaching was performed using a Leica SP8 inverted confocal microscope by scanning the 488 nm and 480 nm beams operating at 100% laser power over a circular region of interest (ROI, diameter 2.4 μm) in the imaging plane, set around the middle of the cell’s height. With an optical section thickness of ~0.85 μm, the cell membrane area that is bleached is approximately a 2.4 μm by 0.85 μm rectangle (in the xz plane). Four frames were acquired at low (2%) laser power for normalization of the fluorescence signal before bleaching the ROI for 13 s as explained above. Recovery was monitored for 30 frames every 1290 ms and then 15-30 frames every min at low laser power (2%). Image size was 512 x 512 pixels. The background was determined with the same ROI area outside the cell using ImageJ and subtracted from the signal.

Postbleach fluorescence recovery traces were fit to F(*t*) = [*F_0_ + F_∞_*(*t*/*t*_1/2_)]/(1 + *t*/*t*_1/2_) where *F_0_* is the initial postbleach fluorescence, *F_∞_* is the asymptote, and *t*_1/2_ is the half-time of recovery. Best-fit values for these parameters were estimated using MatLab’s Curve Fitting Toolbox. The tracer diffusion constant is calculated^70^ as *D_t_* = *A_bleach_*/(4 *t*_1/2_), where *A_bleach_* = 2.4 × 0.85 μm^2^ is the photobleached area. The mobile fraction of proteins is estimated as *F*_∞_ − *F*_0_.

### Tether force as a function of tether extension

Goldfish retinal bipolar neurons were dissociated and plated into Ringers solution without calcium. A 3 μm diameter bead was used to pull a short membrane tether as above. After the tether force stabilized, the tether was extended by moving the cell away from the OT center *via* programmable movement of the piezoelectric stage at 1 μm/s. Images were acquired at a rate of 13-15 frames/s, with *k_Trap_* = 73-75 pN/μm. Changes in tether force or normalized membrane tension vs. extension curves were plotted (Extended Data Figure 2). For chromaffin cells, a similar protocol was used, except a larger trap stiffness (*k_Trap_* = 271 pN/μm) had to be used to prevent the bead from escaping the trap prematurely.

A sudden, short extension of a static tether was used to test the response of the tether force to a step-like tether extension. After a tether was extruded, the tether force was allowed to stabilize for ~10 s. The stage was moved by 1-3 μm at 10-500 μm/s while the force was monitored. Force profiles were normalized to 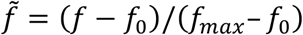 and averaged (Extended Data Figure 2). A double-exponential function was fit to the data using the Matlab Curve Fitting toolbox.

### Fluorescence-based tether force estimates

To test the relationship between the tether force *f_t_* and tether diameter *r_t_*, cells were incubated with 5 μM CellMask Deep Red dye for 15 min in calcium-free Ringers. A short tether was extruded in brightfield mode and held stationary until the force settled. Then the tether was extended at 1 μm/s while fluroescence images were acquired at ~ 4.2 frames/s. Tether force was calculated from the fluorescence channel images as above (CellMask also labeled the beads, allowing their tracking). Tether fluorescence was estimated from a region of interest (ROI, see Extended Data Figure 4 for an example). The number of pixels in the tether ROI was doubled using bicubic interpolation, rotated and the pixels along the direction of the tether were averaged. A Gaussian function was fit to the averaged lateral intensity profile and the area under the Gaussian was taken as the background-corrected tether fluorescence intensity, *I_t_*. This procedure was repeated for every frame in a movie stack. Typical tether diameters are much smaller than the optical sectioning (~0.85 μm); thus, tether fluorescence is integrated over the thickness of the tether. Because 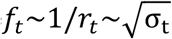 (see above and refs. ^40,62^) and *I_t_*~2π *r_t_l*, where *l* is the length of the ROI along the tether, *f_t_*~1/*I_t_*, and 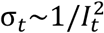. We plotted the relationship between *f_t_* and *I_t_* both in dynamic experiments (where the tether was being pulled at 1 μm/s, e.g. Extended Data Figure 4a,b) and static measurements (where the tether was held at stationary length, Extended Data Figure 4c). In both cases a linear relationship was obtained.

To test how membrane tension changes at the pulling tether are communicated to the probe tether, we pulled two tethers simultaneously as described earlier, but we labeled the cells using CellMask Deep Red as above, and imaged cells in fluorescence (~ 4.2 frame/s). We analyzed the fluorescence of each tether as above, and plotted 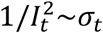 for the two tethers (Extended Data Figure 4d,e).

### Imaging of the F-actin cytoskeleton

Cells were incubated with 1-2 μM of SiR-Actin (Cytoskeleton, Inc. Spirochrome) for 15-30 min in calcium-free buffer, then a tether was pulled as described above. SiR-Actin fluorescence was monitored using 640 nm excitation and a Chroma 485/60 nm, 705/90 nm band-pass emission filter using spinning disc confocal microscopy. We could not detect any Sir-Actin fluorescence in the tethers (5, 5, and 6 tethers extruded from the termini and somata of bipolar cells, and chromaffin cells, respectively). Robust SiR-Actin labeling was evident in every case below the plasma memrbane in intact regions of the cells (Extended Data Figure 3).

### Reagents and materials

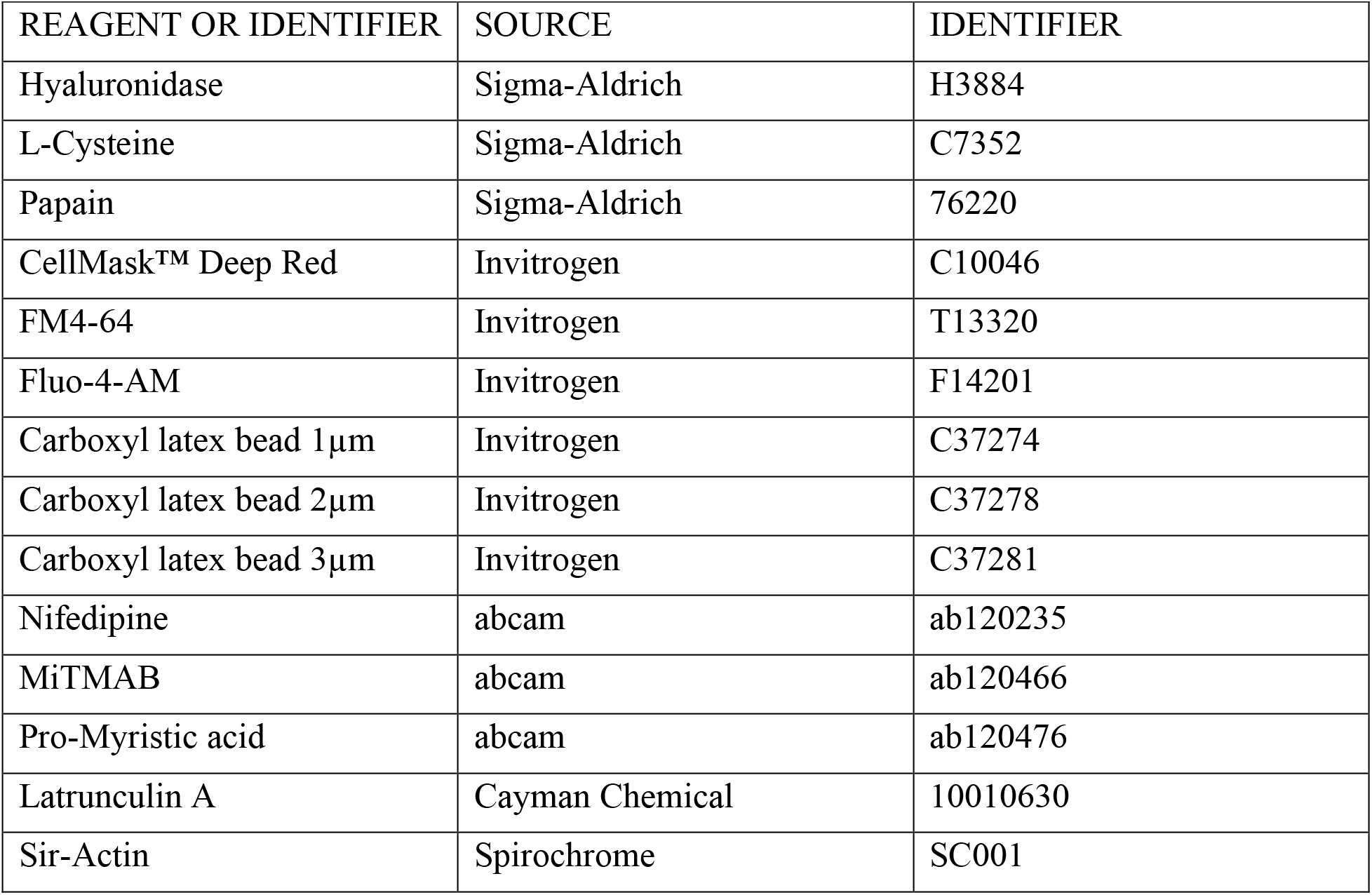

#### Modeling

A detailed description of the numerical simulations and fitting procedures can be found in the Supplementary Information.

#### Statistics

Data were analyzed by MATLAB software. Statistical significance was determined by one-way ANOVA (Extended Data Figure 2c) or Kolmogorov-Smirnov test (CDF comparisons, Extended Data Figure 8).

#### Reporting Summary

Further information on experimental design is available in the Nature Reporting Summary accompanying this paper.

#### Data Availability

The datasets generated during the current study are available from the corresponding author on reasonable request.

#### Code Availability

All code generated to simulate and analyze the data presented in this study is available from the corresponding author on reasonable request.

## Acknowledgements

We acknowledge support from the Yale Kavli Institute for Neuroscience (Innovative Research Award to EK, DZ and BM) and the National Institutes of Health, specifically grants from the National Institute Of Neurological Disorders and Stroke (R01NS122388 and R21NS112754 to EK, DZ and BM) and the National Eye Institute (R01EY032396 to DZ and P30EY026878 Yale Core Grant for Vision Research). We thank members of the Karatekin, Zenisek and Machta labs for discussions, and Pietro De Camilli, Min Wu, and James E. Rothman for critically reading the manuscript.

## Author contributions

NRD and EK conceptualized the initial project with input from DZ. CGP and NRD acquired and analyzed experimental data under the supervision of EK and DZ. MR and BM contributed to modeling and development of quantitative analysis tools. All authors contributed to interpretation of results and design of subsequent experimental, analysis, and modeling steps. EK wrote the initial draft, which was edited and revised by all authors.

## Additional information

Supplementary Information is available for this paper.

Correspondence and requests for materials should be addressed to EK.

---

**Extended Data Figure 1.**
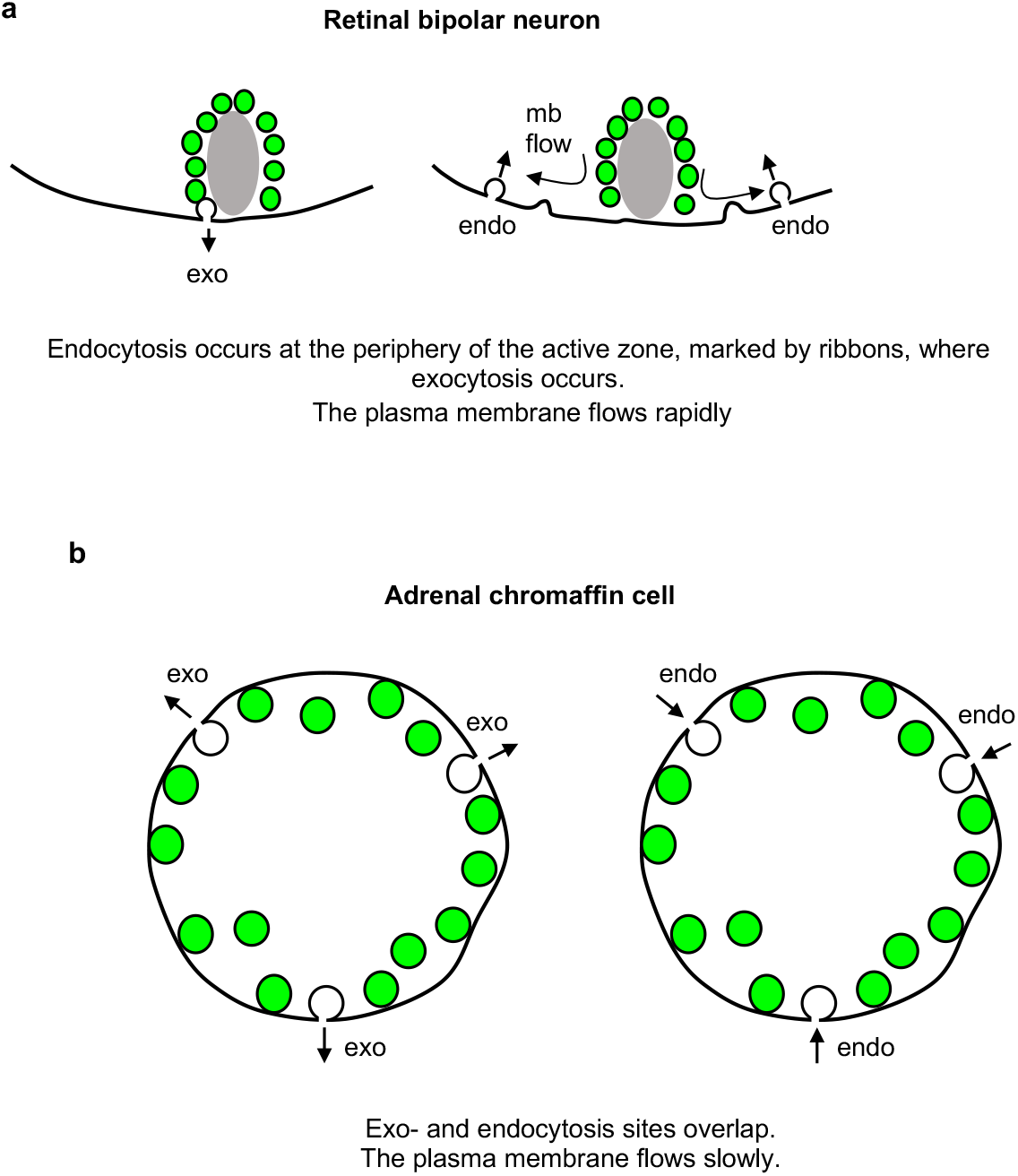
Summary of major findings. **a**. Membrane tension propagates rapidly in the retinal bipolar cell terminal, allowing rapid synaptic vesicle turnover with exo- and endocytosis occurring at distinct loci. **b**. In adrenal chromaffin cells, membrane tension propagates slowly, restricting exo- and endocytosis to occur at overlapping sites.

**Extended Data Figure 2.**
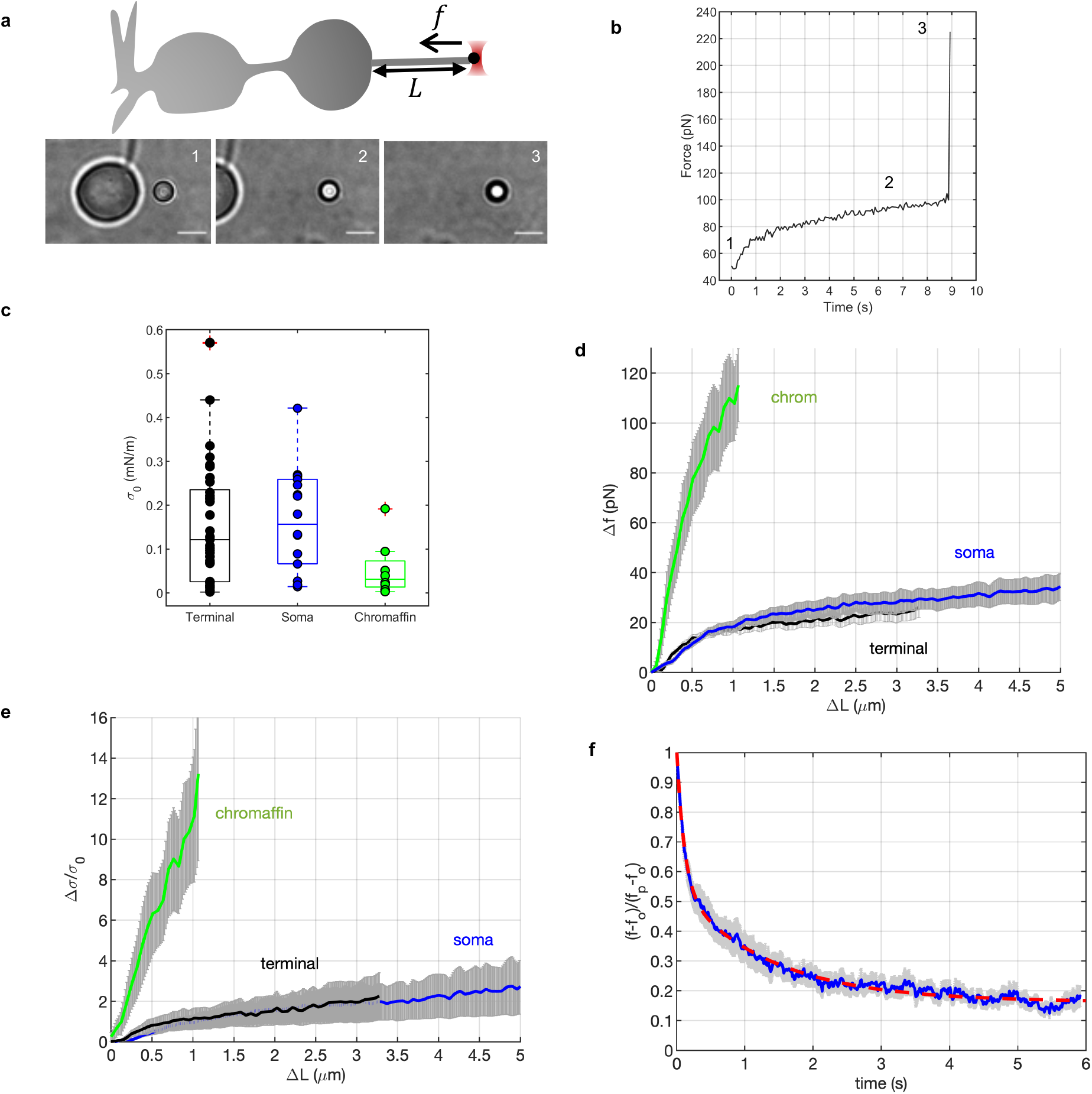
Resting membrane tension and how it changes upon tether extension. **a**. Top: schematic of the experiment. A 3 μm bead held in the optical trap (OT) is used to pull a short, *L*_0_ = 1.5 − 3 μm membrane tether from the cell surface. After the force is stabilized to its static value *f*_0_, the tether is extended at constant speed 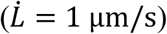 by moving the cell away while the force acting on the bead *f*(*t*) is monitored. Bottom: snapshots of a tether pulled from a bipolar neuronal terminal. Scale bar = 5 μm. **b**. Example of a tether force measurement, for a tether extruded from the terminal for the example shown in **a**. The tether was extended starting *t* = 0 s. The numbers in the snapshots in **a** correspond to the numbers indicated on the force profile. The bead escaped the trap at 3. **c**. Static membrane tension values for bipolar cell terminals (n= 37), somata (n=18), and for chromaffin cells (n=8). There was not a significant difference among the means (p=0.098, 1-way ANOVA). **d**. Change in tether force as a function of tether extension at constant extrusion velocity 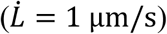 for chromaffin cells (n=6), or somata (n= 8) or terminals (n=9) from bipolar neurons. The gray errorbars represent S.E.M. **e**. The force profiles in **d** replotted as the fractional change in membrane tension (using 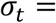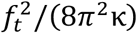, *see Methods*) as a function of extension Δ*L* = *L* − *L*_0_. **f**. Response of the tether force to a sudden extension. A resting tether, drawn from the terminal, was rapidly extended by 1-3 μm. Shown is the average of the resulting changes in the tether forces, rescaled as 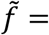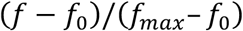. The red dashed line is a fit to 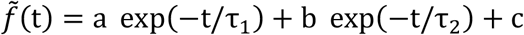, with best fit parameters (with 95% confidence bounds): *a* = 0.47 (0.46, 0.48), *b* = 0.38 (0.38, 0.39), *c* = 0.163 (0.161, 0.164), *τ*_1_ = 0.112 (0.107, 0.116), *τ*_2_ = 1.35 (1.32, 1.39), *R*^2^ = 0.987. (n= 9 tethers.)

**Extended Data Figure 3.**
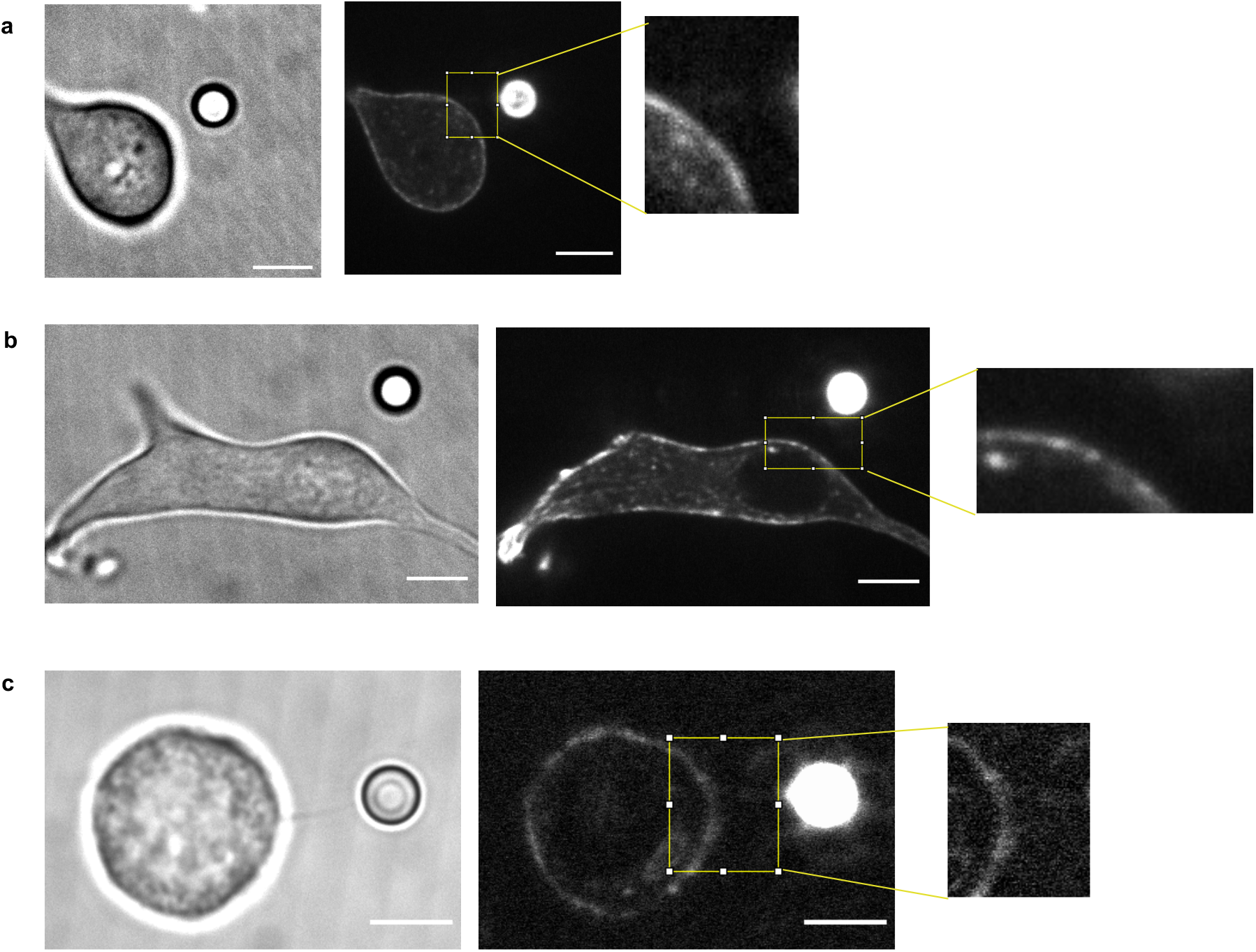
Actin filaments cannot be detected in membrane tethers. **a**. A tether was drawn from the terminal of a bipolar neuron after SiR-Actin labeling of the F-actin cytokeleton. Despite good cortical labeling as reported^43^, no F-actin is detectable in the tether. The experiment was repeated with 4 other cells, with similar results. **b**. A tether drawn from the soma, with no labeling evident in the tether. Two additional cells were tested, with similar results. **c**. A tether drawn from a chromaffin cell, after Sir-Actin labeling. No F-actin can be detected in the tether. Five other cells were tested, with similar results. Scale bars represent 5 μm.

**Extended Data Figure 4.**
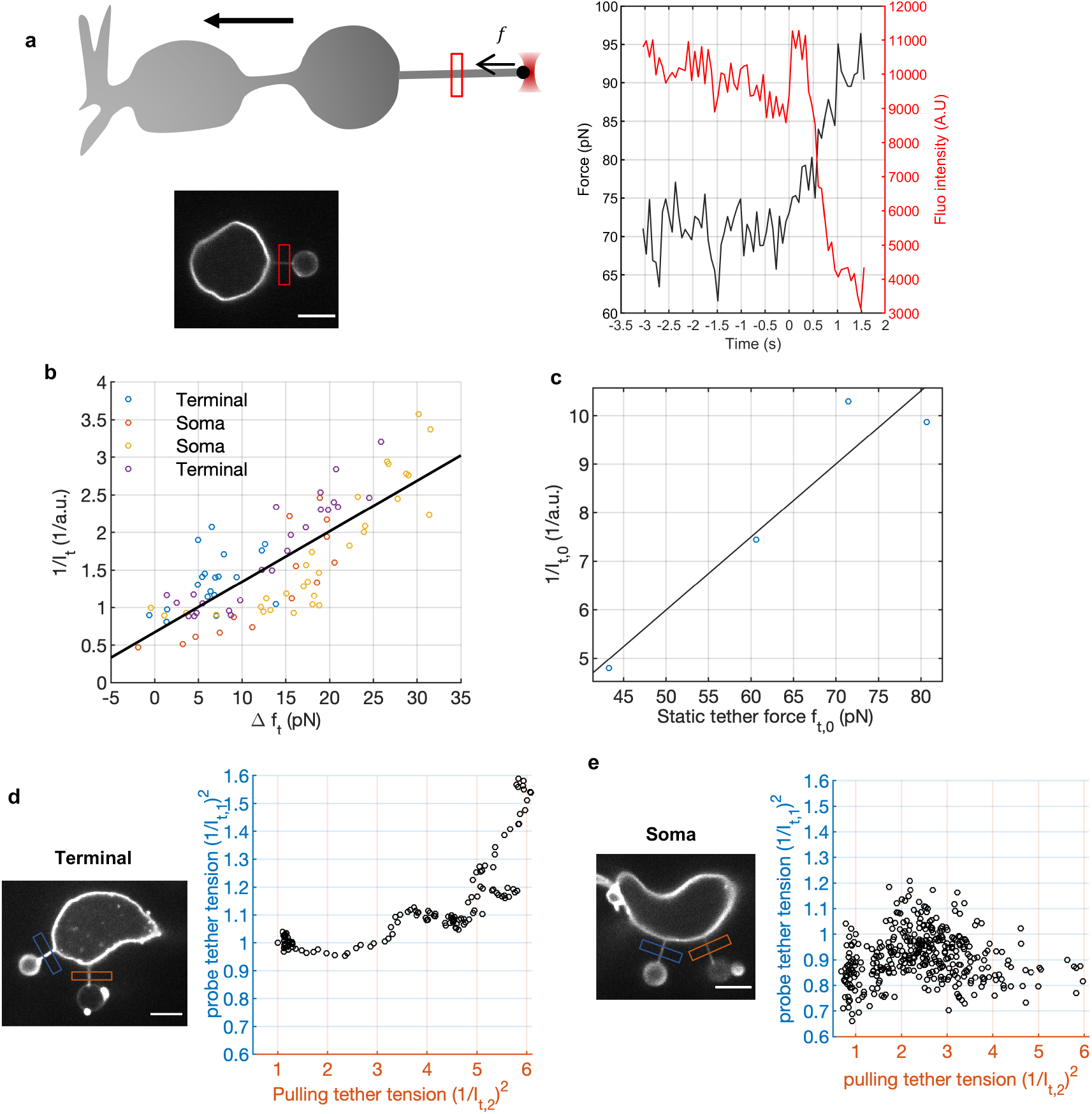
Membrane tension changes tracked using tether fluorescence. **a-c**. Calibration of tether fluorescence as a function of tether force. Cell membranes were labeled with CellMask Deep Red and fluorescence was measured from a fixed area using spinning disc confocal (SDC) microscopy. **a**. Example of a simultaneous force and fluorescence measurement for a tether pulled from the terminal at constant speed (1 μm/s). Left: schematic of the experiment and a snapshot from an image stack recorded while the tether was extended. Right: tether force (left axis, measured using the optical trap) and fluorescence (right axis, from the area marked on the left snapshot) as a function of time. Tether extension started at *t* = 0 s. **b**. Inverse tether fluorescence intensity as a function of the increase in tether force from its initial value. Data from 4 experiments, grouped by color. A linear fit to all data is shown (slope = 6.72 × 10^−6^ AU/pN, *R*^2^ = 0.861). **c**. Relationship between inverse tether fluorescence and tether force for static measurements (e.g. the values for *t* < 0 in **a**). A linear fit had best slope 1.51 × 10^−6^ AU/pN (*R*^2^ = 0.910). **d,e**. Measurement of tether fluorescence simultaneously from two tethers drawn from the same cell. The probe tether (blue box) is held stationary while the pulling tether (red box) is extended at 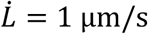. The square of the inverse tether fluorescence intensity ((1/*I_t_*)^2^) is proportional to membrane tension, since the tether force 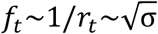 and 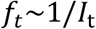 (see b, c). The regions of interest marked with blue (probe tether) and red (pulling tether) boxes were used to calculate the fluorescence intensities (see Methods), which were normalized to their initial values. Scale bars are 5 μm.

**Extended Data Figure 5.**
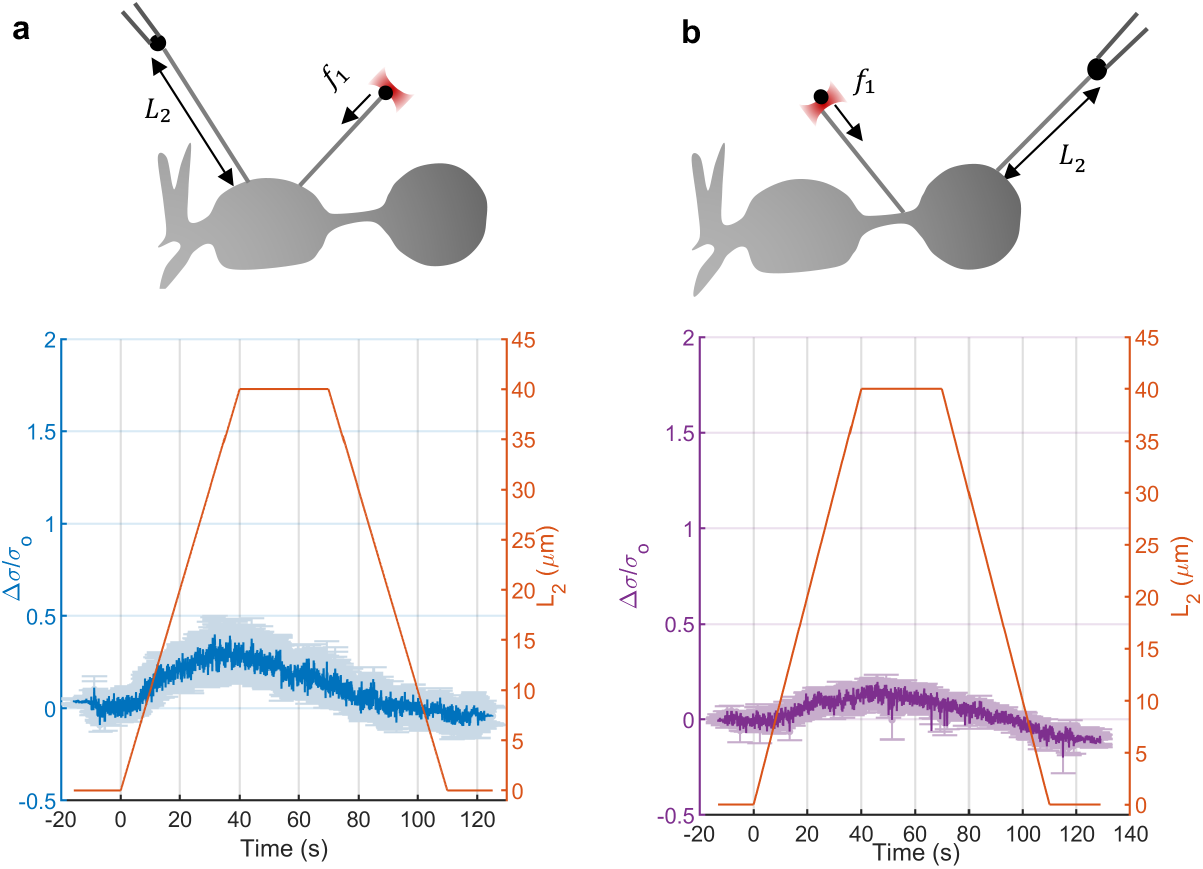
Propagation of membrane tension within the soma and between the terminal and the axon. **a**. Membrane tension change in the probe tether relative to its initial value (left axis) as a function of time as the pulling tether is extended, held stationary, then relaxed (right axis), for tethers pulled from the soma of bipolar neurons (n = 8). Inter-tether distance *d* was 4−12 μm. **b**. Similar measurements, but the pulling tether was extruded from the terminal, while the probe tether was placed in the axon, with inter-tether distances 14-17 μm (n = 4).

**Extended Data Figure 6.**
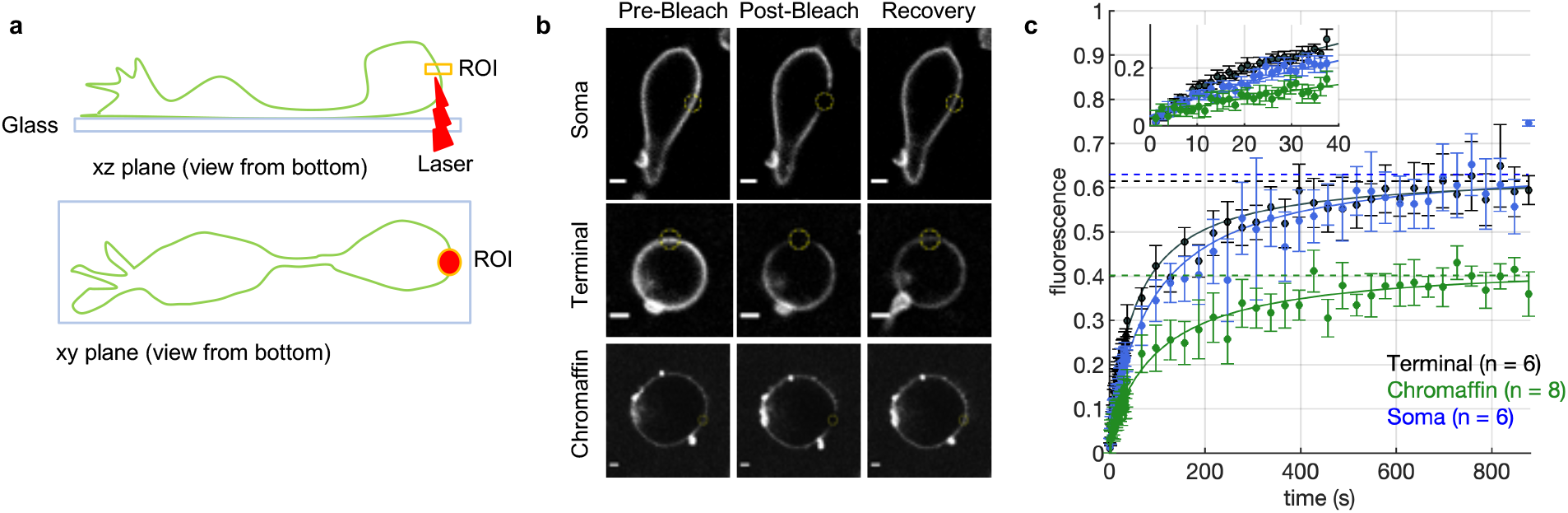
Diffusion and immobile fraction of membrane proteins. **a.** Schematic of the fluorescence recovery after photobleaching (FRAP) measurements for a bipolar cell terminal. Membrane proteins on the cell surface were labeled with Alexa-488 (see Methods). After washing unreacted dye, a 2.4 μm diameter circular region-of-interest (ROI) was bleached in a single optical slice (~0.84 μm thick) in the middle of the terminal. The resulting bleached membrane area is a ~0.84 μm × 2.4 μm rectangle in the xz plane. **b**. Snapshots from actual measurements for a chromaffin cell, a bipolar neuron soma and a terminal (cells are imaged from the bottom). Scale bars are 5 μm. **c.** Fluorescence recovery in the bleached regions was followed as a function of time. The intensity of the ROI for 4 frames prior to bleaching was averaged and used for normalization of the pre-bleach intensity to 1. Recovery was followed initially at 0.773 frames/s for 30 frames, then at a lower rate (0.25-0.50 frames/s) for the remainder of the measurements to minimize photobleaching while capturing the initial rapid phase. Recovery traces for every group were fit to an equation, with the mobile fraction of protein *ϕ_m_* and the tracer diffusion coefficient *D_t_* estimated from the best-fit parameters (see Methods). Best fit values were (with 95% confidence intervals): *D_t_* = (10 ± 7) × 6^−3^, (19 ± 6) × 10^−3^, and (11 ± 5) × 10^−3^ μm^2^/s, and *ϕ_m_* = 0.40 ± 0.03, 0.37 ± 0.02, and 0.37 ± 0.04 for chromaffin cells, termini, and somata, respectively. The dots represent experimental measurements. Inset shows the initial recovery on an expanded scale.

**Extended Data Figure 7.**
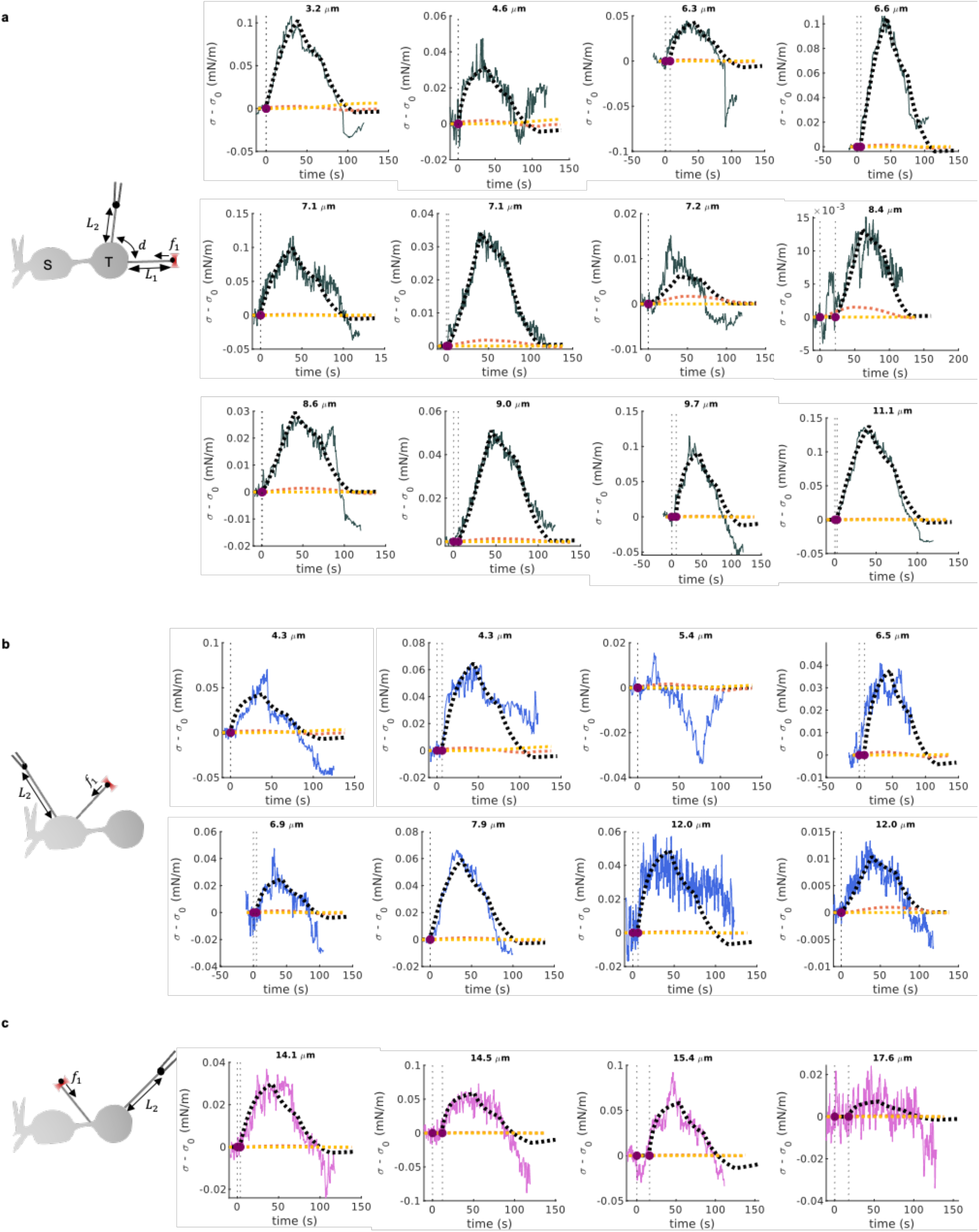
Comparison of experimental measurements of membrane tension propagation with numerical simulations. **a.** A pulling tether was extended from a bipolar cell terminal by 40 μm at 1 μm/s, held for 30 s, then relaxed to its initial extension at 1 μm/s while a probe tether measured the resulting membrane tension changes a distance *d* away. Left, schematic of the experiment. Right: individual measurements of membrane tension changes (black, solid), shown with the predicted tension changes at the probe tether assuming a membrane tension diffusivity *D_σ_* = 0.024 μm^2^/s (yellow, ref. ^16^) or a 100-fold larger value (orange). The model^16^ considers membrane flow through a random array of immobile obstacles (transmembrane domain proteins attached to the underlying cytoskeleton) and predicts diffusive propagation of membrane tension perturbations. The intermembrane distance *d* for every experiment is indicated. Agreement between simulations and measurements is poor in every case. A good match is obtained only assuming *d* = 0.1 μm (black, dashed), suggesting the tension perturbation created by the pulling tether is transmitted with little distance dependence to the probe tether (see Figure 2). **b.** As in **a**, but both tethers were extruded from the soma (blue traces) of bipolar neurons. **c**. As in **a**, but the pulling and probe tethers were extruded from the terminal and the axon, respectively (purple traces).

**Extended Data Figure 8.**
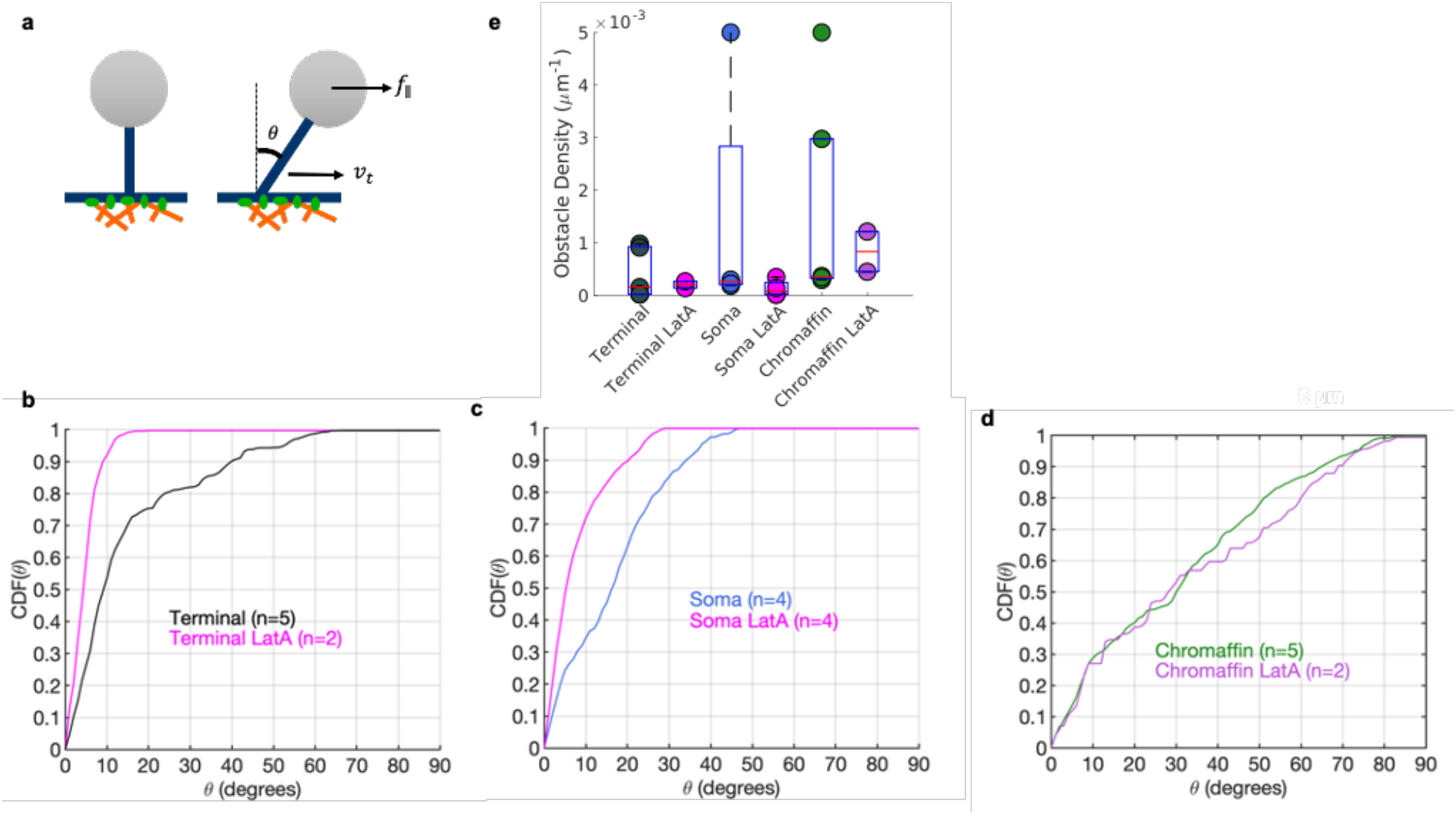
Analysis of membrane tether sliding experiments probing cytoskeleton-membrane friction, related to Figure 3. **a.** Schematic of the experiment. A membrane tether was pulled from the cell surface, then the cell was moved (by moving the xy stage) to create a tangential force *f*_∥_ to drive sliding of the tether’s base with velocity *v_t_* (see Figure 3 and Methods for details). **b**. Cumulative distribution function (CDF) of tether-membrane angles for bipolar cell terminals. Angles were smaller for LatA-treated cells, indicating easier sliding and lower forces. **c**. Similar measurements for tethers drawn from bipolar cell somata. **d**. Same as in c, for chromaffin cells. Angles were smaller for terminals and somata compared to chromaffin cells, indicating easier sliding. LatA treatment shifted *θ* values to lower values in terminals and somata, but not in chromaffin cells (Kolmogorov-Smirnov test, *p* = 7.2 × 10^−12^, 1.6 × 10^−19^, and 6.8.× 10^−5^ for treated vs. untreated terminals, somata, and chromaffin cells, respectively). **e**. Obstacle density for bipolar cell terminals and somata, and chromaffin cells. Obstacles result in tether base stalling (during which *v_t_* < 0.5 μm/s) despite being subject to a substantial tangential force (*f*_∥_ > 5 pN). Obstacle density is defined as the number of individual obstacles encountered during a trajectory, divided by the total distance traveled by the tether base (see Methods).

**Extended Data Figure 9.**
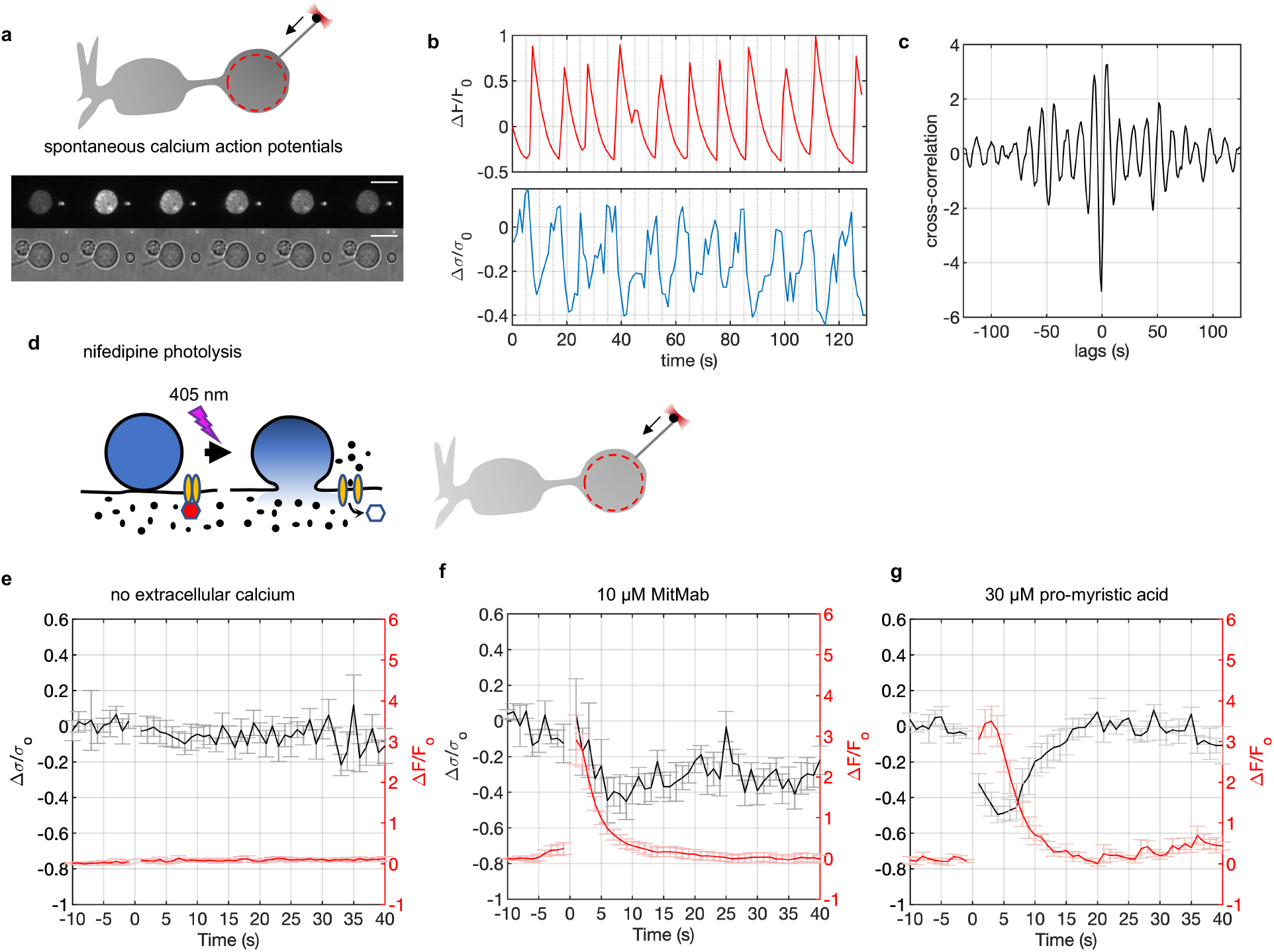
Additional experiments probing the relationship between synaptic vesicle recycling and membrane tension changes. **a**. Top. Schematic of experiments relating spontaneus exo-endocytic activity of bipolar neurons to membrane tension changes in the terminal. A tether was extruded from the terminal using a bead held in an optical trap to monitor membrane tension. The cell was pre-loaded with a fluorescent calcium indicator (Fluo-4) whose fluorescence intensity was monitored in the region of interest (ROI) indicated by the dashed circle in the terminal. Two mM calcium was present in the extracellular medium. Some cells develop spontaneous calcium action potentials (monitored via Fluo-4 fluorescence) that robustly drive exo-endocytosis^36^. Bottom: Snapshots from an image stack of a cell that developed spontaneous calcium activity. Brightfield and fluorescence channels were alternated (1.08 s between pairs of images). Scale bar=5 μm. **b**. Quantification of the Fluo-4 calcium signals and membrane tension measurements for the example shown in **a**. Both the fluorescence (top, red) and membrane tension (bottom, blue) are reported as changes relative to the initial value (Δ*F*/*F*_0_ or Δ*σ*/*σ*_0_). Exocytosis occurs with the upstoke of the calcium wave, followed by endocytosis restoring membrane area^36^. Note that calcium variations anti-correlate with membrane tension changes, consistent with exocytosis lowering and endocytosis restoring membrane tension, respectively. (see Supplementary Movie 6). Five other cells developed spontaneous calcium activity during membrane tension measurements; all such cells also displayed anti-correlated membrane tension variations. **c**. Cross-correlation between calcium signals and membrane tension for data shown in **b**. Notice the strong negative correlation between the two signals at lag ≈ −0.5 s, the time lag between successive fluorescent (calcium) and and bright field (membrane tension) measurements, indicating membrane tension decreases immediately after a calcium increase with our time resolution. **d**. Schematic of the photostimulation protocol (see Figure 4). Calcium signals were integrated from the ROI shown as a dashed red line in the terminal. **e**. When extracellular calcium was omitted, nifedipine photolysis caused changes neither in calcium signals nor in membrane tension. **f.** When endocytosis was inhibited with 10 μM dynamin inhibitor MiTMAB, calcium signals and the initial decrease in membrane tension were not affected, but the recovery of membrane tension was compromised (cf. Figure 4c). **g**. We used pro-myristic acid as a negative control for the MiTMAB experiments. In the cell, pro-myristic acid is rapidly converted to myristic acid which is not a dynamin inhibitor^51^. Membrane tension and calcium dynamics were not appreciably altered compared to controls (Figure 4c). In f and g, 2 mM extracellular calcium was present.

