## Supplementary Discussion for "Rapid propagation of membrane tension at a presynaptic terminal"

**Discussion of modeling work**

**Diffusive model of membrane tension propagation:** It has long been known that modest differences in the immobile fraction of membrane proteins may cause dramatic differences in membrane flow[1,2]. This is predicted qualitatively by the hydrodynamics of flow around a bed of fixed obstacles: in 2-dimensions (2D), obstacles produce long-range perturbations to the flow field that substantially slow bulk flow with increasing obstacle density[2,3], even under conditions when tracer diffusion is relatively unimpeded. A model[3] in which transmembrane domain (TMD) proteins interact with the underlying cytoskeleton and act as a fixed array of obstacles predicts membrane tension propagates diffusively, with a tension diffusion coefficient $D_\sigma = Ek/\eta$, where $E$ is the membrane stretch modulus, $\eta$ is the 2D membrane viscosity, and $k$ is the Darcy permeability of the array of obstacles. Assuming obstacles are randomly distributed, the Darcy permeability is a function of obstacle area fraction $\phi$ and the radius $a$ of obstacles, $k = a^2 f(\phi)$, where $f(\phi)$ is a rapidly-decaying scaling function[4]; for $\phi < 0.37$, $k \approx -\frac{a^2[1+\ln(\phi)]}{8\phi}$. From FRAP measurements of tracer molecules, Shi et al.[3] estimated $\eta \approx 3 \times 10^{-3}$ pN $\cdot$ s/μm and $\phi \approx 0.18$. Combined with published estimates of $E = 40$ pN/μm and $a = 2$ nm, the tension diffusion coefficient was estimated to be $D_\sigma \approx 0.024$ μm$^2$/s for HeLa cells and similarly small for the four other cell types tested. We interpret this as an upper-bound on the speed of tension-diffusion in these cell-types; Shi and coworkers saw no observable tension propagation in HeLa cells over distances of 5 – 15 μm and timescales $\gtrsim 10$ min, which combined with modeling suggest membranes cannot flow faster than what is suggested by this value of $D_\sigma$ (see reference 3, Figure 2)

**Diffusive model cannot explain measured membrane tension dynamics in bipolar neurons:** We used numerical simulations to test if this model could reproduce the membrane tension profiles we measured. We estimated the area fraction of immobile obstacles $\phi$ from fluorescence recovery after photobleaching (FRAP) measurements (Extended Data Figure 6), assuming that the immobile fraction measured in those experiments represents transmembrane proteins that interact with the underlying cytoskeleton. Assuming ~25% of membrane area is occupied by transmembrane proteins[5], we estimate $\phi \approx 0.096 \pm 0.005$ terminal, and $\phi \approx 0.15 \pm 0.005$ in chromaffin cells, yielding $k \approx (6.96 \pm 1.06) \times 10^{-6}$ μm$^2$ and $k \approx (3.06 \pm 2.3) \times 10^{-6}$ μm$^2$ respectively. With these constraints, and the values for $E$, $\eta$, and $a$ above, $D_\sigma \approx 0.093$ μm$^2$/s and 0.048 μm$^2$/s are predicted for the bipolar and chromaffin cells, respectively. Numerical simulations with these values did not produce satisfactory fits to the data for the terminals or somata. Instead, measured profiles more closely followed the predicted tension at the base of the pulling tether, suggesting rapid propagation of membrane tension with a weak distance dependence (Figure 2a, Extended Data Figure 7).

**Tension propagation in bipolar neurons shows no clear distance dependence.** Diffusive mechanisms predict a strong distance dependence in the amplitude and kinetics of a propagating signal. We explored the distance dependence of the kinetics and amplitude of membrane tension perturbations further to determine if the diffusive propagation model proposed by Shi et al.[3] is applicable for bipolar neurons. We plotted the time lag, $t_{sense}$, between the onset of the tension increase at the pulling tether (i.e., when tether extension was initiated) and the onset at the tension increase at the probe tether in Figure 2b. We similarly plotted the maximum change in membrane tension at the probe tether, $\Delta\sigma_{max}$, for a given perturbation created at the pulling tether (Figure 2c). To estimate $t_{sense}$ and $\Delta\sigma_{max}$ accurately and in an unbiased manner, we matched a template to the probe tether profile, by shifting it in time and varying its amplitude until a good match was produced. The template was the model predicted tension profile at the pulling tether shifted in time and rescaled in amplitude (see below for details), which, perhaps surprisingly, yielded curves which provided a good description of the experimental tension profiles at the probe tether. Note, however, that any empirical template that matches the experimental profile well could be used to estimate $t_{sense}$ and $\Delta\sigma_{max}$. We found that $t_{sense}$ and $\Delta\sigma_{max}$ obtained in this manner have no clear distance dependence (Figure 2b,c). The time lag $t_{sense}$ as a function of inter-tether distance could only be well-fit with a diffusion constant of ~24 μm$^2$/s (1000-fold larger than the value estimated[3] for Hela cells) or larger, while $\Delta\sigma_{max}$ could not be fit by simply changing the Darcy permeability $k$ to change $D_\sigma$ (Figure 2b,c). Other membrane parameters may govern tension dynamics. Area expansion modulus measurements are sparse and we take $E = 40\,\text{pN/μm}$, however extreme values up to $E = 1750\,\text{pN/μm}$ have been recorded and suggest substantial cell-to-cell variability[6]. Viscosity measurements are also sparse, but have not been found to vary over orders of magnitude. Overall, these results suggest that in bipolar cells membrane tension propagates rapidly and in a qualitatively different manner from its very slow propagation in chromaffin cells tested here, or other cell types tested by Shi et al.[3]. However, our bipolar cell measurements are consistent with rapid membrane flows reported by Dai and Sheetz[7] during axonal growth.

Although it is difficult to rule out that membrane tension may propagate diffusively in bipolar neurons, a diffusive mechanism would require some combination of the physical parameters (viscosity, stretch modulus, Darcy permeability) to be several orders of magnitude different than those measured in other cell-types, which seems unlikely. Alternative mechanisms could explain rapid membrane flows in bipolar neurons. For example, the model in ref. 3 assumes that obstacles are distributed without particular structure and are immobile during the measurements (that span minutes). If the obstacles are arranged differently, rapid membrane flow would be possible at similar obstacle densities, as proposed by Cohen and Shi[4]. Other possible mechanisms that could contribute to rapid flows are more dynamic connections between the transmembrane domain proteins (the obstacles) and the underlying cytoskeleton[8,9], rapid cytoskeleton turnover, three-dimensional propagation of pressure perturbations[10,11] or other signaling mechanisms. We hope our results will motivate future modeling and experimental work to better understand cell membrane tension propagation mechanisms.

### Numerical methods

**Tension Diffusion Simulations.** Simulations of tension propagation closely followed those detailed in Shi et al.[3]. We simulate the plasma membrane as a disk with a radius $R_{cell} = 100\ \mu m$, divided radially into evenly spaced 0.1 μm concentric circles. We 'pull' an initial tether of length $L_0 = 10$ μm and calculate the tether area $A_t$, tension $\sigma_t$, and radius $r_t$:

$$r_t = \frac{2\pi\kappa}{f_{t,0}} \quad (1)$$

$$\sigma_{t,0} = \frac{\kappa}{2r_t^2} \quad (2)$$

$$A_t = 2\pi r_t L_0 \quad (3)$$

Where κ is the bending modulus of the membrane and $f_{t,0}$ is the initial tether force. We pull or retract the tether at $\dot{L} = 1$ μm/s or hold it at constant length according to the protocol. The tether radius and tension are updated at constant tether area:

$$L \rightarrow L + \dot{L}dt \quad (4)$$

$$r_t \rightarrow \frac{A_t}{2\pi L} \quad (5)$$

$$\sigma_t \rightarrow \sigma_t + \frac{4\pi\kappa L}{A_t^2} \quad (6)$$

Membrane flows on the cell surface with diffusive dynamics, with diffusion coefficient $D_\sigma = Ek/\eta$. The tension across the membrane is determined as solutions as a boundary-value problem PDE for $\sigma(x,t)$, $\frac{\partial\sigma(x,t)}{\partial t} = D_\sigma \frac{\partial^2\sigma(x,t)}{\partial x^2}$, with tension at $\sigma(0,t) = \sigma_t$ and $\sigma(x = R_{cell}, t) = \sigma_0$ between times $t$ and $t + dt$ using MatLab's partial differential equation solver *PDEPE*. Equations were parameterized in axisymmetric coordinates, where $x$ is the radial distance from the tether base. Solutions to these equations at time $t$, $\sigma(x,t)$, were used as initial conditions for the subsequent step's calculation. After updating tension across the cell, lipids flow into or out of the tether. The tether's area change is set by the membrane flux at the cell-tether boundary and the tether tension updated accordingly:

$$\delta A = \frac{2\pi r_t k}{\eta} \nabla_{r=0} dt \quad (7)$$

$$A_t \rightarrow A_t - \delta A \quad (8)$$

$$\sigma_t \rightarrow \sigma_t - \frac{2\pi^2\kappa L^2}{A_t^3}\delta A \quad (9)$$

Solutions were calculated iteratively until the completion of the tether pulling protocol: 10 s at constant tether length, 40 s extension at 1 μm/s, 30 s relaxation at constant tether length, 40 s retraction at 1 μm/s, 30 s relaxation. Solutions can be examined at a given inter-tether distance.

**Extracting $t_{sense}$ and $\Delta\sigma_{max}$ from experiments with the aid of simulations.** Tension measurements from double-tether experiments were smoothed with a Gaussian filter with width set to 1s prior to fitting. We estimated $t_{sense}$, the time to sense at the probe tether a tension perturbation initiated at the pulling tether at $t = 0$, and $\Delta\sigma_{max}$, the maximum change in tension at the probe tether. Individual tension traces were fit to the equation $g(t) = f(t - t_{sense}) \times \Delta\sigma$ where $f(t)$ is the simulated tension at the pulling tether. We fit two parameters, $t_{sense}$ and $\Delta\sigma$, to each experimental curve by minimizing the mean-squared error between the simulation and experimental curve: $mi = \int dt \left(f_{exp}(t) - g(t)\right)^2$.

We extracted $\Delta\sigma_{max}$ from experiments averaging the tension in a half-second window surrounding the maximum recorded tension in the unfiltered trace. $t_{sense}$ and $\Delta\sigma_{max}$ were calculated similarly from simulated traces to give the curves displayed in figures 2b,c. Tension traces retrieved from simulation at inter-tether distances of 0.1 – 20 µm are fit to the simulated tension at the tether base to give $t_{sense}$. $\Delta\sigma_{max}$ was extracted at these inter-tether distances by averaging tension the half-second window of the maximum.


1 Saffman, P. G. & Delbruck, M. Brownian motion in biological membranes. *Proc Natl Acad Sci U S A* **72**, 3111-3113 (1975).

2 Bussell, S. J., Koch, D. L. & Hammer, D. A. Effect of hydrodynamic interactions on the diffusion of integral membrane proteins: tracer diffusion in organelle and reconstituted membranes. *Biophys J* **68**, 1828-1835, doi:10.1016/S0006-3495(95)80359-0 (1995).

3 Shi, Z., Graber, Z. T., Baumgart, T., Stone, H. A. & Cohen, A. E. Cell Membranes Resist Flow. *Cell* **175**, 1769-1779 e1713, doi:10.1016/j.cell.2018.09.054 (2018).

4 Cohen, A. E. & Shi, Z. Do Cell Membranes Flow Like Honey or Jiggle Like Jello? *Bioessays* **42**, e1900142, doi:10.1002/bies.201900142 (2020).

5 Dupuy, A. D. & Engelman, D. M. Protein area occupancy at the center of the red blood cell membrane. *Proc Natl Acad Sci U S A* **105**, 2848-2852, doi:10.1073/pnas.0712379105 (2008).

6 Needham, D. & Hochmuth, R. M. A Sensitive Measure of Surface Stress in the Resting Neutrophil. *Biophysical Journal* **61**, 1664-1670, doi:Doi 10.1016/S0006-3495(92)81970-7 (1992).

7 Dai, J. & Sheetz, M. P. Axon membrane flows from the growth cone to the cell body. *Cell* **83**, 693-701, doi:10.1016/0092-8674(95)90182-5 (1995).

8 Sens, P. Rigidity sensing by stochastic sliding friction. *Epl-Europhys Lett* **104**, doi:Artn 38003 10.1209/0295-5075/104/38003 (2013).

9 Brochard-Wyart, F., Borghi, N., Cuvelier, D. & Nassoy, P. Hydrodynamic narrowing of tubes extruded from cells. *Proc Natl Acad Sci U S A* **103**, 7660-7663, doi:10.1073/pnas.0602012103 (2006).

10 Charras, G. T., Yarrow, J. C., Horton, M. A., Mahadevan, L. & Mitchison, T. J. Non-equilibration of hydrostatic pressure in blebbing cells. *Nature* **435**, 365-369, doi:10.1038/nature03550 (2005).

11 Moeendarbary, E. *et al.* The cytoplasm of living cells behaves as a poroelastic material. *Nature Materials* **12**, 253-261, doi:10.1038/nmat3517 (2013).
