## Supplementary material for "Rapid propagation of membrane tension at a presynaptic terminal": SI Guide, captions of SI Movies

The following supplementary materials accompany the manuscript:

**SI_Discussion_210526.docx**
A detailed description of the numerical simulations and fitting procedures can be found in the Supplementary Information Appendix.

**Supplementary Movie 1.** Related to Figure 1c. Testing propagation of membrane tension in a bipolar neuronal terminal using a double-tether protocol. Frames were acquired at ~33 Hz. Scale bar is 5 μm.

**Supplementary Movie 2.** Related to Figure 1e. Testing propagation of membrane tension in a chromaffin cell using a double-tether protocol. Frames were acquired at ~17 Hz. Scale bar is 5 μm.

**Supplementary Movie 3.** Image stack showing sliding of a membrane tether drawn from the terminal of a bipolar neuronal terminal. The cell membrane was labeled with the lipophilic dye FM4-64. Elements of the tracking, the bead position, position of the tether- base, membrane contour, and tether-membrane tangent line, are overlayed in magenta. Images collected at 8.77 frames/s. The tether slides freely along the membrane. Related to Figure 3.

**Supplementary Movie 4.** Image stack showing attempt to drag a membrane tether drawn from a chromaffin cell whose membrane was labeled with CellMask Deep Red. Elements of the tracking, the bead position, position of the tether base, membrane contour, and tether-membrane tangent line, are overlayed in magenta. Images collected at 4.34 frames/s. The tether did no slide along the membrane despite application of tangential forces. Related to Figure 3.

**Supplementary Movie 5.** Image stack showing tether sliding in a retinal bipolar neuron soma. The membrane was labeled with CellMask Deep Red. Elements of the tracking, the bead position, position of the tether base, membrane contour, and tether-membrane tangent line, are overlayed in magenta. Images collected at 5.55 frames/s. The tether slides briefly after a tangential force is applied, but becomes stuck again. Related to Figure 3.

**Supplementary Movie 6.** Related to Extended Data Figure 9a-c. Image stack showing spontaneous intracellular calcium variations anti-correlate with membrane tension changes in a bipolar neuronal terminal. Two channels were alternated during acquisition: brightfield (to detect bead position for membrane tension estimation, 30 ms exposure) and fluorescence (to detect calcium variations using Fluo-4, excited at 488 nm, 200 ms exposure). They are overlaid for presentation purposes. The period between a pair of images was 1.08 s. Scale bar is 5 μm.
